# Coagulative Granular Hydrogels with an Enzyme Catalyzed Fibrin Network for Endogenous Tissue Regeneration

**DOI:** 10.1101/2024.11.20.624078

**Authors:** Zhipeng Deng, Camila B. Tovani, Simona Bianco, Gianni Comandini, Aya Elghajiji, Dave J. Adams, Fabrizio Scarpa, Michael R. Whitehouse, Annela M. Seddon, James P. K. Armstrong

## Abstract

Granular hydrogels, composed of densely packed microgels, are an emerging class of injectable microporous scaffolds that provide interstitial porosity for endogenous cell recruitment and tissue repair. However, weak bonding interactions between constituent microgels compromises the mechanical integrity of these biomaterials, limiting their scope and effectiveness for *in vivo* application where structural support is required. To address this challenge, we introduce a new bioinspired stabilization method and a novel class of regenerative biomaterial: *coagulative granular hydrogels*, assembled from thrombin-functionalized gelatin methacryloyl microgels. The surface-bound thrombin is enzymatically active and catalyzes the conversion of fibrinogen into a fibrin hydrogel that extends throughout the interstitial voids of the granular hydrogel. This secondary network acts as a biological glue to stabilize the granular hydrogel, yielding shear and compressive properties comparable to bulk hydrogel controls. Furthermore, the interstitial fibrin network provides a favorable microenvironment for the adhesion, proliferation, and invasion of endothelial cells, highlighting the potential of the biomaterial to support endogenous tissue repair. This innovative stabilization mechanism provides a responsive biomaterial system, and future studies will seek to explore how the biomaterial is annealed *in vivo* by the presence of blood plasma fibrinogen.

## 1. Introduction

Although the human body has a capacity for self-healing, tissue damage is not always fully or correctly resolved to allow normal function. This has led to the development of regenerative biomaterials, engineered with biophysical and biochemical cues that can stimulate, accelerate, or guide endogenous tissue repair.^[1]^ Polymer hydrogels are an attractive class of regenerative biomaterials providing highly hydrated three-dimensional (3D) networks that can mimic native extracellular matrices and provide supportive and inductive environments for tissue repair.^[2]^ However, the use of bulk hydrogels with continuous crosslinked polymer networks can present a number of challenges. For instance, bulk hydrogels are not generally injectable, and while it is possible to design systems that undergo sol-gel transitions *in vivo*,^[3]^ this approach greatly constrains the choice of biomaterial. In addition, the dense nanoscale structure in bulk hydrogel networks can substantially limit host cell infiltration and nutrient mass transport.^[4]^

These limitations have been addressed by an emerging class of biomaterials known as granular hydrogels, which consist of densely packed (jammed) microgels. These injectable biomaterials retain the advantageous properties of bulk hydrogels (*e.g.*, hydration, biochemical cues, nanoscale topography), while also providing an interstitial microporous network to allow the infiltration of endogenous cells and nutrients.^[5,6]^ These properties have driven extensive exploration of granular hydrogels as injectable porous biomaterials for regenerative medicine and tissue repair.^[7]^ However, a major limitation of granular hydrogels is their substantially inferior mechanical properties compared to bulk hydrogels, which is due to weak bonding interactions between the constituent microgels^[8]^. Consequently, granular hydrogels can be readily broken down by dynamic mechanical loading and can even undergo passive degradation under static conditions.^[9]^

Many approaches have sought to stabilize granular hydrogels by strengthening the interparticle interactions. An attractive option is reversable binding mechanisms, such as hydrazone bonding, electrostatic interactions, and host-guest chemistries,^[10–13]^ which can preserve injectability while also stabilizing the static state. These biomaterials, however, still fall short of matching the bulk hydrogel mechanics. Greater stabilization can be achieved through the formation of covalent bonds *via* photo-annealing,^[14–18]^ however, these crosslinking methods limit matrix remodeling and translation as an injectable biomaterial.^[19]^ Covalent bonds can also be formed by enzyme mediated annealing, for example, through the addition of factor XIII to catalyze the crosslinking of peptide-functionalized granular hydrogels prior to *in vivo* injection.^[20–22]^ It should be noted that all these cited studies use granular hydrogels with empty interstitial voids, which can be filled by a secondary hydrogel network.^[23–27]^ These composite biomaterials have greatly improved mechanical performance but have compromised injectability or require post-curing of the secondary network, both factors that can restrict clinical translation.

This highlights the unmet need for a method that can preserve the injectability of granular hydrogels while matching the bulk hydrogel mechanics. Here, we introduce a new bio-inspired stabilization method and a novel class of regenerative biomaterial, namely *coagulative granular hydrogels* composed of thrombin-functionalized microgels (**Figure 1a**). Thrombin is an important enzyme in wound healing that catalyzes the assembly of soluble blood plasma fibrinogen into crosslinked fibrin networks.^[28]^ The coagulative granular hydrogels harness this enzymatic reaction to catalyze the conversion of fibrinogen into a secondary fibrin network that fills the interstitial voids (**Figure 1b**). We have designed a robust fabrication method to produce coagulative GelMA granular hydrogels and have comprehensively assessed their enzymatic activity, composite structure, shear/compressive mechanics, and interactions with endothelial cells and spheroids. We have demonstrated that the fibrin network stabilizes the granular hydrogel, with the resulting composite providing compressive/shear mechanics on par with bulk GelMA hydrogel. Moreover, this fibrin hydrogel mimics the initial stage of wound healing, providing the granular hydrogel with a conducive microenvironment for cell adhesion and proliferation.

**Figure 1.**
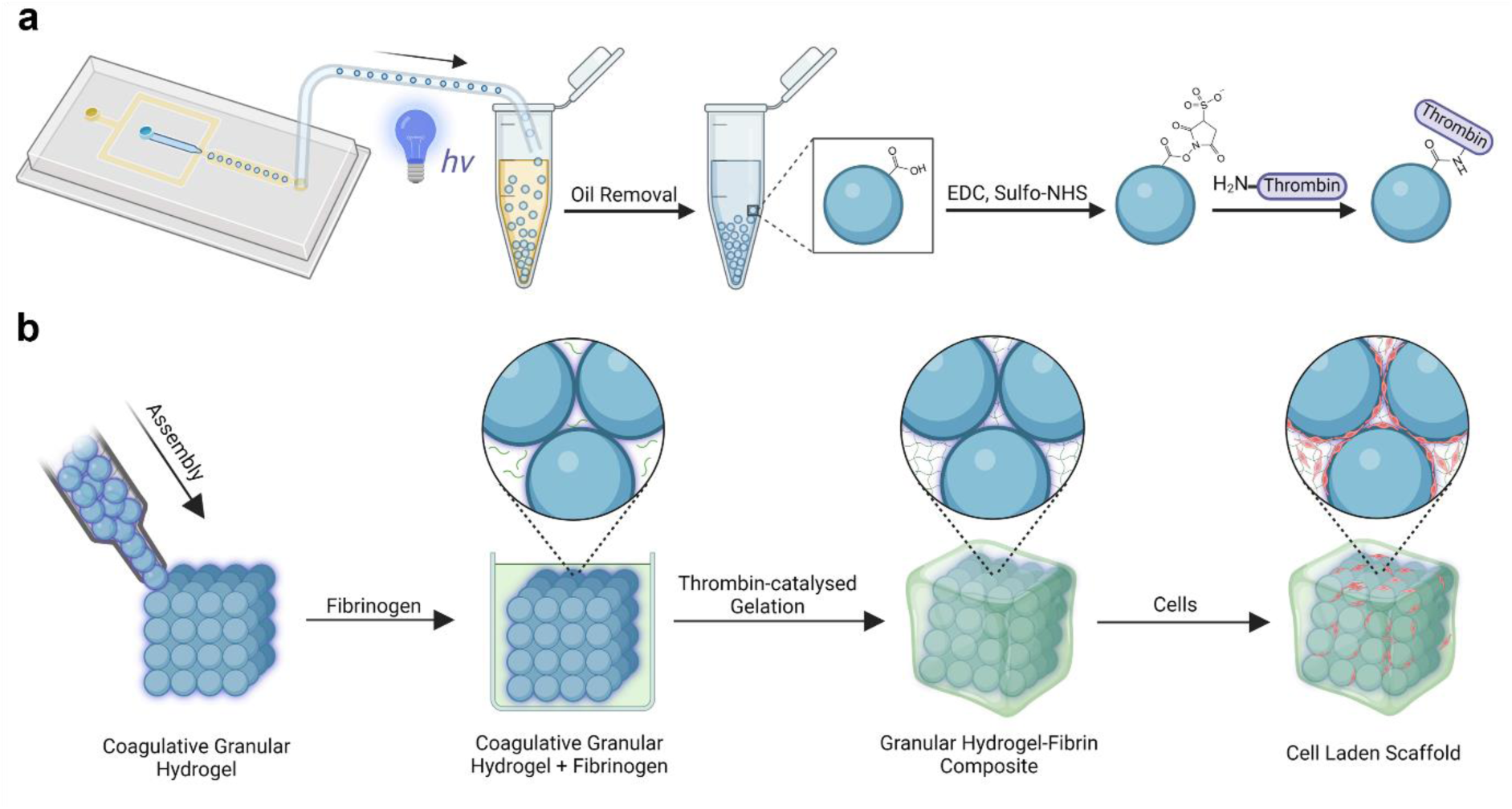
The fabrication and testing of coagulative granular hydrogel scaffolds. a) GelMA microgels are fabricated using emulsion microfluidics, transferred into an aqueous phase, and functionalized with thrombin using a 1-ethyl-3-(3-dimethylaminopropyl)carbodiimide (EDC) / *N*-hydroxysulfosuccinimide (sulfo-NHS) reaction. (b) Thrombin-functionalized microgels are assembled into a coagulative granular hydrogel, which is then cultured with fibrinogen and endothelial cells to model how the biomaterial might support fibrin formation and cell growth *in vivo*.

Overall, coagulative granular hydrogels overcome the limitations of other granular hydrogel systems, providing a method to stabilize the biomaterial to the level of the bulk hydrogel without compromising injectability. Importantly, the stabilization mechanism is responsive to an *in vivo* wound healing environment, which will allow future clinical applications to trigger fibrin network formation through exposure to blood plasma fibrinogen. The carbodiimide chemistry used to generate the thrombin-functionalized microgels is generic and should be readily applied to other granular hydrogels or other enzymes in the wound healing cascade. By addressing key limitations in both bulk and granular hydrogels, this versatile approach paves the way for *in vivo* application across a variety of endogenous tissue regeneration scenarios.

## 2. Results and Discussion

### 2.1. Fabrication and Characterization of Thrombin-functionalized Microgels

We used an established protocol to synthesize GelMA using gelatin and methacrylic anhydride.^[29]^ Proton nuclear magnetic resonance (^1^H NMR) spectroscopy was used to confirm the substitution of the free gelatin amines (**Figure S1**, Supporting Information),^[30]^ while a fluoraldehyde assay confirmed a consistent degree of functionalization of 91 ± 2% across the five batches used in this study (**Figure S2**, Supporting Information). We designed and made a microfluidic chip with a flow-focusing geometry (**Figure S3**, Supporting Information), which we used to generate size-tunable microdroplets containing an aqueous phase of 8% w/v GelMA solution surrounded by a continuous phase of surfactant-stabilized mineral oil. The inclusion of 8.5 mM lithium phenyl (2,4,6-trimethylbenzoyl) phosphinate (LAP) in the aqueous polymer phase enabled efficient photocrosslinking of the microdroplets into GelMA microgels, which were washed and then phase transferred into phosphate buffered saline (PBS). The crosslinked GelMA microgels were spherical, with a diameter of 160 ± 14 µm in oil, which increased to 200 ± 16 µm in PBS due to the swelling of polymer chains in aqueous solution (**Figure 2a-b**).

**Figure 2.**
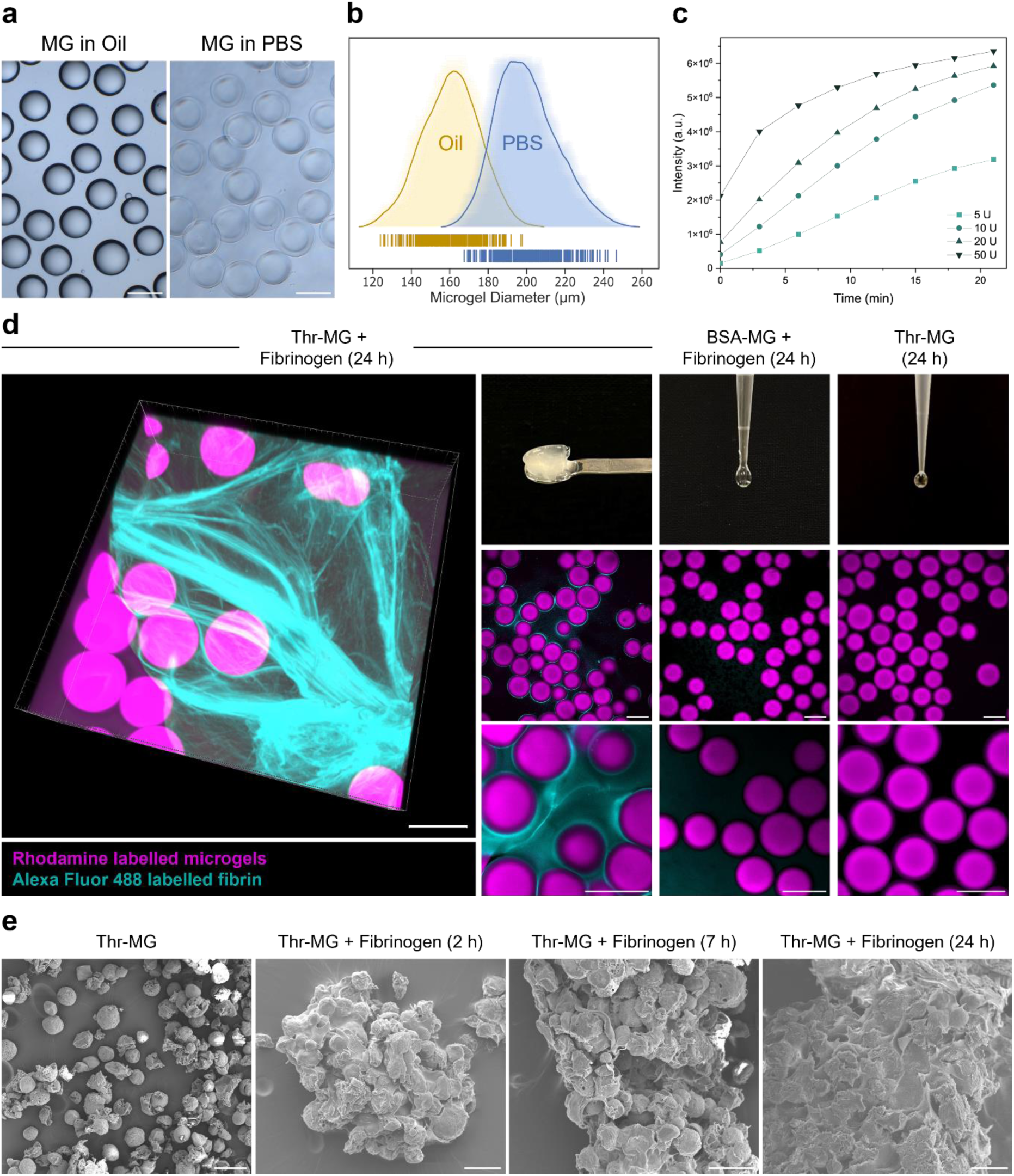
Characterization of thrombin-functionalized GelMA microgels (Thr-MG). a) Representative brightfield and phase contrast images of microgels in mineral oil and PBS. Scale bar: 200 µm. b) Average shifted histogram of the microgel diameter in mineral oil (160 ± 14 µm) and PBS (200 ± 16 µm), as measured from 317 microgels. c) Enzymatic activity of Thr-MGs with increasing thrombin used in the reaction (5, 10, 20, 50 U). d) Confocal fluorescence microscopy and camera images of Thr-MG after 24 h of fibrinogen incubation, as well as controls of BSA-MG with fibrinogen and Thr-MG without fibrinogen. The microscopy study was performed using rhodamine-functionalized microgels (magenta) and Alexa Flour 488 conjugated fibrinogen (cyan). Scale bar: 200 µm. e) SEM images of Thr-MGs alone, and after 2, 7, and 24 h of incubation with fibrinogen. Scale bar: 100 µm.

We used carbodiimide-based chemistry to immobilize thrombin on the surface of the GelMA microgels. The activity of the surface-immobilized thrombin was evaluated using a 7-amino-4-methylcoumarin fluorometric assay (**Figure 2c**). Enzymatic activity was detected in the thrombin-functionalized microgels (Thr-MG) but not in the surrounding buffer, indicating a successful coupling reaction (**Figure S4a**, Supporting Information). When 5 U of thrombin was used, 37% of the catalytic activity was preserved after crosslinking. This value was similar to controls in which thrombin was incubated overnight without any microgels or crosslinking reagents (43% activity) (**Figure S4b**, Supporting Information). This thrombin concentration, which was used for all subsequent experiments, yielded Thr-MGs with an enzymatic activity of ∼0.11 U mg^−1^ (by dry mass). It should be noted that the enzymatic activity of the microgels could be increased by raising the concentration of thrombin during the reaction, providing tuneability for different applications. Importantly, there was negligible enzymatic activity in both control groups: unfunctionalized microgels (MG) and microgels functionalized with bovine serum albumin (BSA-MG).

We next sought to investigate the interaction between Thr-MG and the fibrinogen substrate at 37℃. Brightfield microscopy showed that fibrinogen induced the aggregation of Thr-MG after just 15 min (**Figure S5,** Supporting Information) to form a stable gel after 24 h (**Figure 2d**). High magnification brightfield images revealed fibers surrounding the Thr-MGs (**Figure S6**, Supporting Information), which suggested that the soluble fibrinogen had been successfully catalyzed into a secondary fibrin network that effectively “glued” the microgels together. Interestingly, microgel controls without thrombin also formed loose aggregates following the addition of fibrinogen but did not form a gel after 24 h. These results indicated that fibrinogen molecules induced the early aggregation of microgels, while the catalysis of fibrinogen into fibrin was mediated by the surface-immobilized thrombin. We next used confocal fluorescence microscopy to visualize fluorescently-labelled Thr-MGs exposed for 24 h to a fluorescently-conjugated fibrinogen. These images showed bright halos of fibrin around the Thr-MGs, while 3D rendering clearly revealed a dense fibrillar network surrounding and bridging the microgels (**Figure 2d** and **Video S1**, Supporting Information). This network was present throughout the full volume of the Thr-MG sample but was absent in the control groups (BSA-MGs with fibrinogen, Thr-MG without fibrinogen).

Scanning electron microscopy (SEM) was used to examine the development of the fibrin gel at different timepoints after addition of fibrinogen (2, 7, 24 h) **(Figure 2e**). The Thr-MGs were initially discrete microgels with smooth surfaces and open pores, but 2 h of incubation with fibrinogen produced small aggregates covered in fibers. After 7 and 24 h, these fibers had assembled into larger strands and bundles, connecting the microgels to form a densely packed fibrillar network. High magnification images confirmed that the fibril morphology in the Thr-MG samples closely matched a bulk fibrin gel control (**Figure S7**, Supporting Information). In contrast, no fibers were observed in the control groups without thrombin functionalization (**Figure S8**, Supporting Information). Taken together, the confocal fluorescence microscopy and SEM images provided strong evidence that the formation of fibrin fibers was specifically catalyzed by the surface-immobilized thrombin.

### 2.2. Coagulative Granular Hydrogel Scaffold

Having verified the enzymatic activity of the Thr-MGs, we used centrifugation to form jammed granular hydrogels (Thr-GH). The granular hydrogel could be readily extruded by hand through a 26-gauge needle (inner diameter = 0.26 mm) to form long, continuous strands (**Figure 3a**). Brightfield microscopy of the extruded Thr-GH filaments confirmed that the individual microgels remained intact. These results are consistent with studies that use granular hydrogels as injectable biomaterials that flow at high strain and then rapidly recover into an elastic solid at low strain.^[10,31–33]^ Shear thinning was confirmed by rheology, which was used to measure a sharp decrease in the viscosity of Thr-GH with increasing shear rate (**Figure 3b**). The samples were next subjected to cycles of low and high strain in order to assess shear recovery (**Figure 3c**). The Thr-GH behaved like an elastic solid at 0.1% strain, with a storage modulus (G′ = 21.9 ± 2.9 kPa) substantially higher than the loss modulus (G′′ = 0.8 ± 0.1 kPa). The Thr-GH rapidly transitioned to a liquid-like state at 200% strain (G′ = 0 kPa, G′′ = 0.1 ± 0.0 kPa) and then quickly recovered when returned to 0.1% strain (first cycle G′ = 25.3 ± 6.3 kPa, G′′ = 1.0 ± 0.3 kPa; second cycle G′ = 25.3 ± 6.0 kPa, G′′ = 1.0 ± 0.4 kPa). This non-Newtonian behavior, dependent on interparticle interaction and friction,^[6,15,32]^ matched the unfunctionalized granular hydrogel (GH) controls (**Figure S9**, Supporting Information), which confirmed that the shear thinning and recovery were not affected by thrombin functionalization.

**Figure 3.**
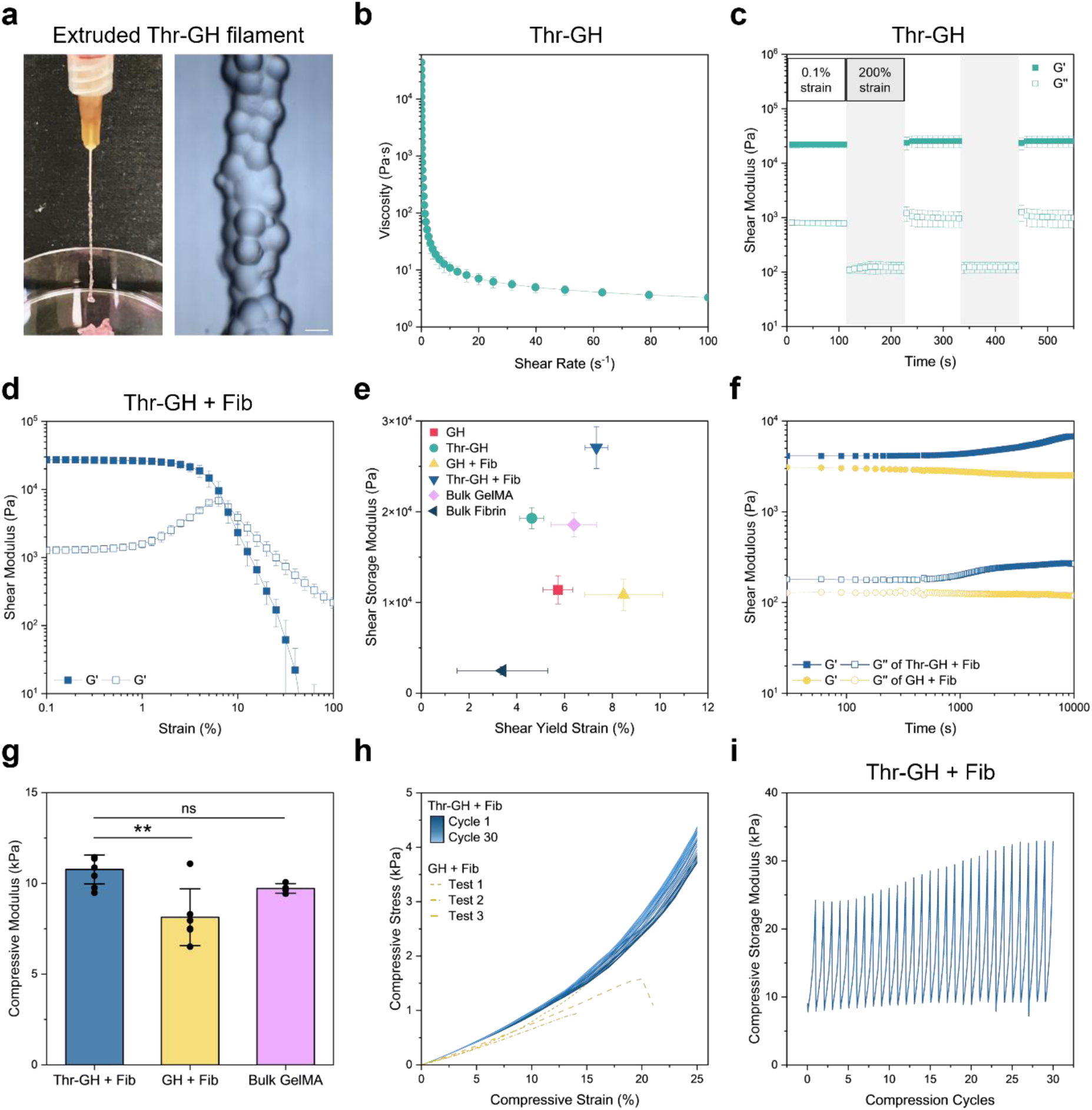
Coagulative granular hydrogel rheology and compression testing. a) Camera image and representative brightfield image of a thrombin-functionalized granular hydrogel (Thr-GH) manually extruded through a 26-gauge needle. Scale bar: 200 µm. b) Viscosity measurement of Thr-GH with shear rates from 0-100 s^−1^. c) Shear recovery test of Thr-GH, measuring shear storage (G’) and loss moduli (G’’) at alternating 0.1% strain (unshaded) and 200% strain (shaded). d) Strain sweep of Thr-GH-fibrin composite (Thr-GH + Fib), measuring G’ and G’’ across 0.1-100% shear strain. e) Comparison of G’ and yield strain of Thr-GH + Fib and controls. f) Time-resolved rheology showing the change in G’ and G’’ after addition of fibrinogen to GH or Thr-GH. g) Characterization of the dynamic compressive moduli of Thr-GH + Fib, GH+ Fib and a bulk GelMA hydrogel, calculated as the slope between 2-7% strain. h) Cyclic compression tests of Thr-GH + Fib under dynamic cyclic loading from 0-25% strain for 30 cycles. The stress strain curves of GH + Fib samples failed to reach 25% strain on the first cycle (shown as yellow dashed line). i) Change in the compressive storage modulus (E’) of Thr-GH + Fib over 30 dynamic compression cycles. For viscosity, shear recovery test, strain sweeps, storage moduli, yield strains, and compressive moduli, data are presented as mean ± standard deviation, with a sample size of n ≥ 3. Statistical analysis performed using Kruskal-Wallis test with Dunn’s post-hoc test. **p < 0.01, ns = no significance (p > 0.05).

The Thr-GH samples incubated with fibrinogen for 24 h (Thr-GH + Fib) could be handled readily and were highly stable against disaggregation. This stability was visually demonstrated by imaging granular hydrogels subjected to continuous agitation. While the Thr-GH control disintegrated after 15 min of agitation and completely dissociated after 2 h, the Thr-GH + Fib composite remained stable throughout (**Figure S10**, Supporting Information). Rheology was also used to quantify the dynamic shear mechanical properties of the Thr-GH-fibrin composite (Thr-GH + Fib) with granular and bulk hydrogel controls. The profile of the strain/frequency sweeps were consistent with the literature,^[15,34,35]^ presenting linear trends in G′ at low frequency as well as broad linear viscoelastic regions with strain-independent G′ (**Figure 3d, Figure S11-12**, Supporting Information). While the yield strain was similar for all samples (between 5 and 8%), the G′ of Thr-GH + Fib (27.1 ± 2.3 kPa) was significantly higher than the five controls: Thr-GH (19.3 ± 1.2 kPa), GH + Fib (10.8 ± 1.7 kPa), GH (11.4 ± 1.6 kPa), bulk GelMA hydrogels (18.6 ± 1.3 kPa), and bulk fibrin hydrogels (2.5 ± 0.4 kPa) (**Figure 3e**). These results indicated that the secondary fibrin network throughout Thr-GH dissipates stress under shear, leading to an improved shear modulus that exceeded even the bulk hydrogel controls. We also used time-resolved rheology to monitor the temporal changes in the granular hydrogels during incubation with fibrinogen (**Figure 3f**). Thr-GH showed increases in G′ and G′′ after 500 s that eventually plateaued at ∼10000 s, while the shear moduli values of the GH control remained stable throughout.

While these rheology studies confirmed the response to shear, it was important to understand how the biomaterial would respond to compressive loads that might be present *in vivo*.^[36]^ Dynamic compression tests were performed under wet conditions to mimic physiological conditions.^[37]^ The compressive modulus of Thr-GH + Fib (10.8 ± 0.8 kPa) was significantly higher than the GH + Fib control (8.1 ± 1.6 kPa) and, notably, was comparable to the bulk GelMA hydrogel control (9.7 ± 0.3 kPa) (**Figure 3g**). We next sought to repeatedly compress the granular hydrogels to 25% strain to understand the biomaterial response to cyclic loading. The GH + Fib controls all disintegrated at the first loading cycle, with failure measured between 14-21% strain. In contrast, we successfully compressed the Thr-GH + Fib composite to 25% strain over 30 cycles without failure (**Figure 3h**). This repeated loading led to a gradual ∼30% increase in compressive storage modulus (**Figure 3i**) and a ∼15% decrease in sample thickness (**Figure S13**, Supporting Information). Collectively, these results indicated that the Thr-GH + Fib composite behaved elastically and could endure repeated compression at high strain with load-induced stiffening attributed to an increase in microgel packing density of the granular hydrogel phase.

The microgel packing density is a critical property of granular hydrogels that significantly impacts mechanical properties as well as the ability to support cell and vessel infiltration.^[38]^ Volumetric reconstruction of light sheet fluorescence microscopy images (**Figure 4a** and **Figure 14**, Supporting Information) was used to calculate a microgel packing density of 0.67 for Thr-GH. This value suggested that the deformable nature of the microgels enabled a packing density midway between the theoretical maxima for hard spheres in random close packed (0.64) and cubic/hexagonal close packed (0.74) configurations.^[39]^ Interestingly, the Thr-GH + Fib composite had a lower microgel packing density (0.54), with the interstitial space filled by fibrin. This finding indicated that the formation of fibrin fibers pushes apart the microgels slightly, thus increasing the void fraction compared to Thr-GH alone. This expansion provides a new approach to increase the void fraction without compromising the structural integrity and mechanical properties of the granular hydrogels.

**Figure 4.**
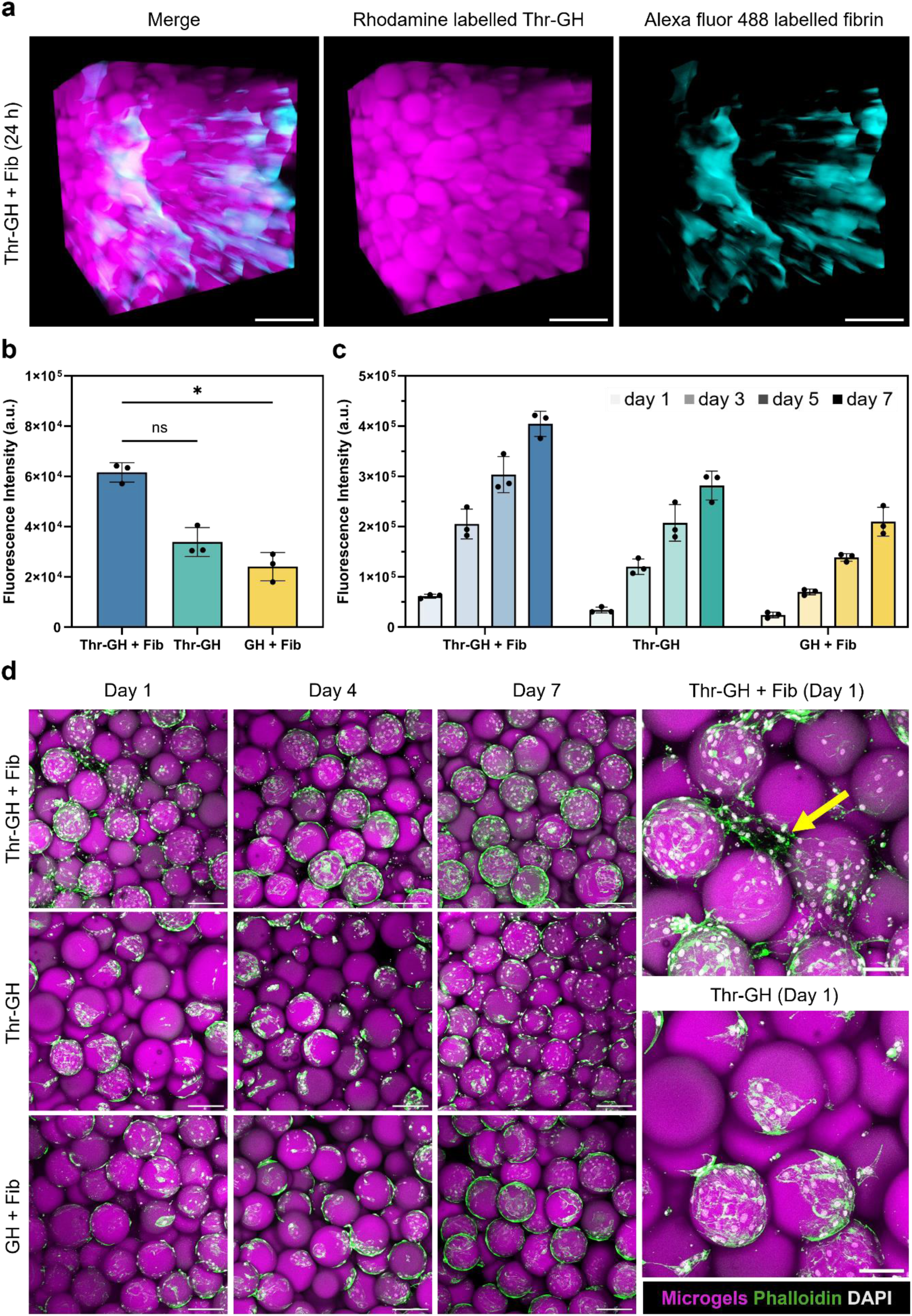
Volumetric imaging and *in vitro* cell culture. a) 3D reconstructed stacks of Thr-GH + Fib using light sheet fluorescence microscopy. Thr-GHs were made using GelMA that was fluorescently labelled with rhodamine (shown in magenta) and incubated with Alexa Flour 488 conjugated fibrinogen (shown in cyan). Scale bar: 500 µm. b) Quantification of cell metabolism in Thr-GH + Fib, Thr-GH and GH + Fib on Day 1. Data are presented as mean ± standard deviation, with a sample size of n ≥ 3. Statistical analysis performed using Kruskal-Wallis test with Dunn’s post-hoc test. *p < 0.05, ns = no significance (p > 0.05). c) Quantification of cell metabolism in Thr-GH + Fib, Thr-GH and GH + Fib over 7 days. d) Representative maximum intensity projection images of HUVECs cultured on Thr-GH + Fib, Thr-GH, GH + Fib over 7 days. HUVECs exhibited spreading in the interstitial fibrin network of Thr-GH + Fib (indicated with yellow arrow), which was not seen in the Thr-GH control. Granular hydrogels were made using GelMA that was fluorescently labelled with rhodamine (shown in magenta), while the cells were stained for F-actin filaments (phalloidin, shown in green) and nuclei (DAPI, shown in white). Scale bar: 200 µm.

### 2.3. In Vitro Cell Culture

The interstitial fibrin network was designed not only to stabilize the granular hydrogel but also to support endogenous cell regeneration. We thus performed a series of *in vitro* cell studies to model whether the Thr-GH + Fib composite would support the adhesion and proliferation of endothelial stem cells. We seeded human umbilical vein endothelial cells (HUVECs) on Thr-GH + Fib and compared the cell response to granular hydrogel controls. An alamarBlue assay performed on day 1 revealed a higher metabolic activity for the cells on Thr-GH + Fib compared to the Thr-GH and GH + Fib controls (**Figure 4b**), strongly suggesting that the fibrin matrix aids cell adhesion to the biomaterial. All samples supported cell proliferation, with Thr-GH + Fib maintaining a substantially higher metabolic activity than the controls on days 3, 5 and 7 (**Figure 4c** and **Figure S15**, Supporting Information). To visualize the endothelial cell morphology within the granular hydrogels, samples were fixed on days 1, 4 and 7 of culture, stained for F-actin and DNA, and then imaged using multiphoton microscopy (**Figure 4d**). Qualitatively, more cells were present on the Thr-GH + Fib samples than the controls, which tallies with the observation from the alamarBlue study that the fibrin network aided cell adhesion. While all samples supported cell proliferation, there were notable differences in the location and morphology of the cells. In particular, the Thr-GH + Fib samples exhibited endothelial cells spreading throughout the interstitial fibrin network, whereas the cells on the granular hydrogel controls were bound exclusively to the microgel surface. Collectively, these *in vitro* results demonstrate the enhanced capabilities of Thr-GH + Fib in supporting endothelial cell adhesion and proliferation, highlighting the potential of this biomaterial for endogenous tissue repair.

We next sought to model how the presence of interstitial fibrin would support the formation and morphology of sprouting vascular networks. We generated multicellular vascular spheroids containing a mixture of HUVECs and human mesenchymal stem cells (hMSCs). These vascular spheroids were embedded within Thr-GH, which was then treated with fibrinogen and cultured for 3 d with vascular endothelial growth factor stimulation for the final 2 d. Confocal microscopy performed at day 3 showed that the Thr-GH + Fib composite effectively supported vascular sprouting and invasion from the spheroids into the interstitial fibrin network (**Figure 5a**). We observed substantially less vascular sprouting without the addition of fibrinogen (*i.e.*, the Thr-GH control), with the spheroid cells primarily remaining adhered to the surface of the microgels. The size of the vascular network was compared to a day 0 spheroid, which revealed substantial horizontal and vertical outgrowth for the Thr-GH + Fib composite (5.6 ± 0.2 fold; 2.8 ± 0.1 fold, respectively). These values were significantly greater than the corresponding outgrowth observed for Thr-GH (3.6 ± 0.2 fold; 1.6 ± 0.4 fold, respectively) (**Figure 5b**). This difference in vascular outgrowth was visualized by re-slicing the volumetric images (**Figure 5c**). Together, these findings underscore the crucial role of the interstitial fibrin in facilitating the invasion of a vascular network into the granular hydrogel. This emphasizes the potential of the coagulative granular hydrogels to effectively support cellular integration and promote endogenous tissue repair.

**Figure 5.**
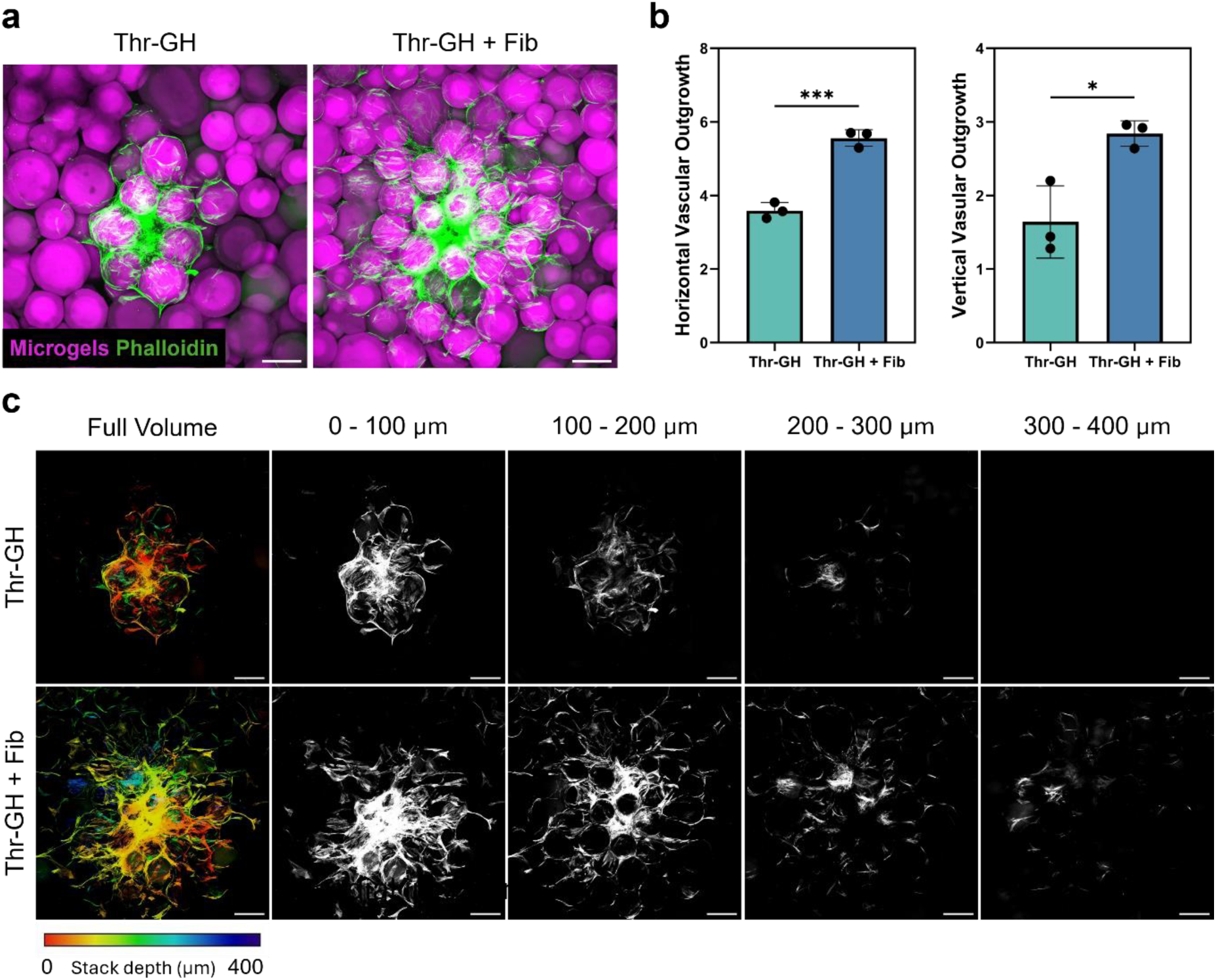
Vascular spheroid sprouting assay. a) Representative maximum intensity projection images of HUVEC-hMSC spheroidscultured within Thr-GHand Thr-GH + Fib for 3 days. Thr-GHs were made using fluorescently labelled GelMA (shown in magenta), while the spheroids were stained for F-actin filaments (phalloidin, shown in green). Scale bar: 200 µm. b) Quantification of the horizontal and vertical outgrowth in Thr-GH and Thr-GH + Fib, with the average radii of the day 3 vascular networks normalized to a day 0 spheroid. Data are presented as mean ± standard deviation, with a sample size of n = 3. Statistical analysis was performed using unpaired t test with Welch’s correction. ***p < 0.001, *p < 0.05. c) Representative z-projections of vascular spheroid sprouting in Thr-GH and Thr-GH + Fib, generated from phalloidin-stained samples. The full volume is shown color-coded by z-height while 100 µm thick sections are shown in greyscale at different depths through the samples. Scale bar: 200 µm.

## 3. Conclusions

In this study, we have developed an injectable coagulative granular hydrogel using GelMA microgels functionalized with thrombin. We demonstrated that surface-immobilized thrombin can effectively catalyze fibrinogen into fibrin, resulted in the *in situ* formation of a secondary fibrin network. The fibrin network acts to “glue” the microgels together, with the granular hydrogel / fibrin composite exhibiting shear and compressive mechanical properties exceeding the bulk GelMA hydrogels. Furthermore, the microporous granular structure and the interstitial fibrin network was shown to support endothelial cell adhesion, proliferation, and invasion. Overall, the properties of these coagulative granular hydrogels are highly aligned with the goals of endogenous tissue regeneration: *i.e.*, an injectable biomaterial that would be mechanically reinforced *in vivo* by the presence of blood plasma fibrinogen to form a stable and supportive microenvironment for host cell invasion. Although these results need to be validated *in vivo*, this study provides a generalizable design feature that can be widely applied to injectable granular hydrogels designed for endogenous tissue regeneration.

## 4. Experimental Section

### Gelatin Methacryloyl (GelMA) Synthesis

GelMA was synthesized as previously reported.^[29]^ Type A gelatin from porcine skin (300 g Bloom, Merck G1890) was dissolved to 100 mg mL^−1^ in ultrapure water (18.2 MΩ cm) under stirring at 50°C. Methacrylic anhydride (Merck 276685) was then added to the solution at a ratio of 0.6 mL per gram of gelatin. The solution was stirred vigorously for 3 h and then centrifuged at 3500 g for 3 min at room temperature. The supernatant was collected, diluted with 50 mL of pre-warmed ultrapure water (∼40°C), and then transferred into a 12-14 kDa regenerated cellulose dialysis tubing (Fisher Scientific 11465859). The solution was dialyzed against 5 L ultrapure water at 40°C for 6 d with the water changed twice a day. After dialysis, the pH of the solution was adjusted to 7.4 with 1 M sodium bicarbonate (VWR RC-091) and sterile-filtered with a Corning vacuum filter system (0.22 µm, Scientific Laboratory Supplies 430769). The GelMA solution was then freeze-dried using the Benchtop Pro 3L ES-S5 freeze dryer (SP Scientific) and stored at −20°C for later use.

### GelMA Characterization

^1^H-NMR spectroscopy was performed to confirm the conjugation of methacrylate groups. 20 mg of GelMA and gelatin were dissolved separately in 0.75 mL deuterium oxide (VMR 1.13366). 700 µL of each sample was loaded into NMR tubes (Norell NORS55007), with ^1^H-NMR spectra obtained using a Jeol ECZ 400 MHz NMR spectrometer. Fluoraldehyde™ *o*-Phthaldialdehyde Reagent Solution (Fisher Scientific 10530054) was used to quantify the degree of functionalization of each GelMA batch. Gelatin standards of 1, 0.5, 0.2, 0.1, 0.05, 0.02 mg mL^−1^ were prepared in PBS (Fisher Scientific 10388739). Each batch of GelMA was dissolved in PBS to a concentration of 1 mg mL^−1^. 300 µL of each sample/standard were mixed with 600 µL fluoraldehyde reagent solution. Upon mixing, 250 µL triplicates of each sample/standard mix were transferred to individual wells of a black 96-well plate (Greiner Bio-One 655090). Fluorescence was measured (λ_ex_ = 360 nm, λ_em_ = 450 nm) using Synergy Neo2 microplate reader (BioTek).

### Microfluidic Chip Fabrication

Molds for T-junction microfluidic chips were designed on Solidworks^®^ (Dassault Systèmes) and printed using a Form 3B+ SLA 3D printer (Formlabs). After removing the printing supports, the 3D printed molds were cured at 60°C for 30 min. Polydimethylsiloxane (PDMS) was made by mixing silicone elastomer base and curing agent from Silicone Sylgard 184 Kit (Scientific Laboratory Supplies 63416.5S) at a 10:1 mass ratio, which were then poured into the 3D printed molds and baked at 60°C for 4 h. Fully cured PDMS chips were removed from the mold and cleaned with ethanol. The PDMS chips were plasma treated using a FEMTO low-pressure plasma cleaner (Diener Electronics) for 30 s under 100 W and 1 bar O_2_, and then immediately sealed with a plasma-treated chip base to create water-tight microchannels. This assembly was then baked in the oven at 60°C for 1 h to enhance sealing between the PDMS and the base. The chip design can be found in the Supplementary Data (**Figure S3**, Supporting Information).

### Microgel Fabrication

A 5 mL Terumo syringe containing a solution of 80 mg mL^−1^ uncrosslinked GelMA and 2.5 mg mL^−1^ LAP (Merck 900889) in PBS was loaded onto an AL1010 syringe pump (World Precision Instruments). A second syringe containing mineral oil (Merck M8410) with 5% v/v Span^®^ 80 (Merck S6760) was loaded onto an identical syringe pump. The GelMA/LAP flow rate and the mineral oil flow rate were set to 3 and 30 µL min^−1^, respectively. The two solutions were flowed into the microfluidic chip *via* low-density polyethylene tubing (Fisher Scientific 13190873). Once a continuous flow of monodisperse microdroplets was observed by eye in the outlet tubing, the microdroplets were crosslinked in one of two ways. *In situ crosslinking* was performed inside the outlet tubing with a UV lamp (365 nm, ∼19 mW cm^−2^, Vilber), with the resulting microgels centrifuged (2000 g, 2 min, room temperature) and washed with 0.1% v/v Tween-20 (Fisher Scientific 10066100) in PBS and then PBS alone to remove the oil phase before storing at 4°C. *Crosslinking after washing* was performed by transferring uncrosslinked microdroplets to 15 mL falcon tubes, which were cooled to −20°C, then centrifuged at 2000 g for 2 min. The supernatant was removed and replaced with mineral oil. The washing process was repeated, with the microd roplets resuspended first in mineral oil, then in PBS, and finally in PBS with 2.5 mg mL^−1^ LAP. The microdroplets were crosslinked in a trough on an ice bath for 1 h under the UV lamp. The microgels were collected and washed with PBS a further 3 times before storing at 4°C. The latter protocol was used for the gelation test (brightfield microscopy and SEM), while the former protocol was used for all other experiments.

### Fluorescent Microgel Fabrication

GelMA was dissolved in PBS to a concentration of 20 mg mL^−1^. NHS-rhodamine (Thermo Fisher Scientific 46406) was added to the solution at a ratio of 30 mg per gram of GelMA. The solution was stirred vigorously for 5 h at 50°C in the dark. After the reaction, the solution was transferred into a 12-14 kDa regenerated cellulose dialysis tubing and dialyzed against 5 L ultrapure water at 40°C for 6 d with water changed twice a day. After dialysis, the pH of the solution was adjusted to 7.4 with 1 M sodium bicarbonate and sterile-filtered with 0.22 µm Corning vacuum filter system. The rhodamine-tagged GelMA was freeze-dried and stored at −20 °C for later use. Fluorescent GelMA microgels were prepared and functionalized as above but using a 1:9 mixture of rhodamine-GelMA and GelMA instead of pure GelMA.

### Size Analysis

The diameter of microgels in oil and PBS were measured using ImageJ (Fiji) (n *>* 300). The threshold of the images was adjusted to reveal the outlines of the microgels, then the ‘analyze particle’ function was used to measure the Feret’s diameter of the microgels. The size distributions were plotted as average shifted histograms as described elsewhere.^[40]^

### Thrombin Functionalization

Microgels were centrifuged at 2000 g for 2 min, with the pellet re-suspended in 0.1 M (2-(*N*-morpholino)ethanesulfonic acid) (MES) buffer at pH 5.5 (Fisher Scientific 15442958). This process was repeated four times, before a 250 µL pellet (equivalent to 20 mg of freeze-dried microgels) was suspended in 3 mL of MES buffer. 1 mL of 40 mg mL^− 1^ EDC (Merck 341006) and 1 mL of 65 mg mL^−1^ sulfo-NHS (Merck 56485) were added to the reaction. The suspension was stirred at room temperature for 30 min, before removing the supernatant. The activated microgels were then resuspended in 3 mL of PBS and 50 µL of 100 U mL^−1^ thrombin (Merck T1063) and stirred at room temperature overnight. After the reaction, thrombin-functionalized microgels (Thr-MG) were washed with PBS (5 times, 2 min, 2000 g) to remove any unbound thrombin and reaction byproducts. Microgel controls were prepared as above but without the addition of thrombin (EDC-activated microgels, EDC-MG) or with the addition of 1 mL of 10 mg mL^−1^ bovine serum albumin (Merck A2153) in PBS instead of thrombin (BSA-functionalized microgels, BSA-MG). All microgels were stored in PBS at 4 °C until further use.

### Thrombin Activity Assay

The enzymatic activity of the Thr-MG, the microgel controls, and the supernatant from the functionalization process was measured using a thrombin activity fluorometric assay kit (Merck MAK242), performed according to the manufacturer’s protocol. Briefly, 50 µL of each diluted sample was added to a white 96-well plate (Greiner Bio-One 655207), alongside 50 µL thrombin standards (0, 5, 10, 15, 20, 25, 50, 100 ng per well). Each sample/standard was mixed with 50 µL of substrate mix. Fluorescence signals were measured (λ_ex_ = 350 nm, λ_em_ = 450 nm) every 5 min for 60 min using a microplate reader at 37℃. Based on the plotted data, two time points from the linear regions of samples and standards were chosen to calculate the difference in fluorescence (ΔRFU) of each. The standard curve was plotted as concentration of thrombin vs ΔRFU, which was used to determine the thrombin concentration of the samples.

### Imaging of Microgel-Fibrin Network

Fibrinogen from human plasma (Thermo Fisher Scientific F3879) was dissolved in phenol-free Dulbecco’s Modified Eagle Medium/Nutrient Mixture F-12 (DMEM/F-12, Thermo Fisher Scientific 21041025) to a concentration of 50 mg mL^−1^. 150 µL of either Thr-MG, EDC-MG, or BSA-MG pellets were resuspended in 400 µL of DMEM/F-12 (equivalent to a microgel concentration of 30 mg mL^−1^) and then mixed with 250 µL of fibrinogen solution in 4-well plate (Thermo Fisher Scientific 176740). A fibrin gel control was prepared the same way by mixing 400 µL of a 2.5 U mL^−1^ thrombin solution with 250 µL of fibrinogen solution. The well plate was sealed with parafilm and placed on an orbital shaker at 50 rpm and 37℃. The morphology of the microgel samples were observed at different reaction intervals (0, 2, 7, 24 h) using brightfield microscopy (MOTIC S3). These samples were also fixed for SEM using 2.5% v/v glutaraldehyde in 0.1 M sodium cacodylate buffer (pH 7.4) for 2 h. The samples were rinsed twice with ultrapure water and then dehydrated by 5 min immersions in 30, 50, 70, 80, 90, 99.8, 100% v/v ethanol. Dehydrated samples were dried using a Leica CPD300 critical point dryer, then sputter coated with gold (∼12.5 nm) using a Quorum Emitech K575X metal sputter coater at 40 mA sputter current for 60 s before imaging using FEI Quanta 200 scanning electron microscope. Confocal fluorescence microscopy was also used to visualize the microgel-fibrin network. Fluorescent Thr-MGs and BSA-MGs were resuspended in DMEM/F-12. 50 µL of each suspension was mixed with 100 µL of a 20 mg mL^−1^ fluorescent fibrinogen solution in separate wells of an 8-well uncoated µ-slide (Thistle Scientific 80801). Fluorescent fibrinogen solution was prepared by dissolving 5 mg of Alexa Fluor 488-tagged fibrinogen (Thermo Fisher Scientific F13191) and 45 mg of fibrinogen in 2.5 mL PBS. After incubation at 37℃ overnight, the samples were imaged using multi-laser confocal laser scanning microscope (Leica SP5II and Leica SP8), equipped with a 488 nm argon laser and 561 nm solid state yellow laser. Images were acquired using 10X and 20X dry objective lenses with a step size of 4.28 µm used for 3D stacks. Images were exported using LAS X software (Leica), processed using ImageJ (Fiji) and reconstructed in 3D using ImarisViewer.

### Formation of Jammed Granular Hydrogels

Jammed granular hydrogels (GH, Thr-GH) were made by centrifuging microgel suspensions at 2000 g for 2 min in either 15 mL Falcon tubes using an Eppendorf centrifuge 5702 or a 48-well plate using a SIGMA 4-16K centrifuge. After centrifugation, the supernatant was removed.

### Imaging of Composite Granular Hydrogels

Thr-GH made from rhodamine-tagged Thr-MG were transferred to cryomolds (Agar Scientific AGG4581) before the addition of either 20 mg mL^−1^ Alexa Fluor 488-tagged fibrinogen solution in PBS or PBS alone (no fibrinogen control). After overnight incubation at 37℃, the two samples (Thr-GH ± Fib) were removed from the mold and glued onto a sample holder. The sample holder was inserted into the imaging chamber of a Zeiss Z.1 light sheet fluorescence microscope. The imaging chamber was filled with PBS and images were acquired with a z-step size of 1.23 µm using 488 and 561 nm argon lasers and 5X dry objective lens. The images were 3D rendered using arivis Vision4D software. The void area in the granular hydrogels was segmented from the acquired image stacks. Packing density was calculated as the total stack volume minus the void fraction, all divided by total stack volume.

### Injectability Test

Thr-GH and unfunctionalized granular hydrogels (GH) were loaded into 2.5 mL Terumo syringes fitted with 26-gauge needles (Fisher Scientific 15301557) and extruded manually. An extruded sample was prepared on a glass coverslip and imaged using brightfield microscopy.

### Rheology

All rheological measurements (n = 3) were performed using Anton Parr Physica MCR301 rheometer equipped with a 12.5 mm diameter parallel plate geometry. Measurements were performed with samples loaded in a sandblasted 24-well plate (Greiner Bio-One 662160) attached to the plate at 37℃. Four granular hydrogel groups were tested (Thr-GH ± fibrinogen, GH ± fibrinogen), these were centrifuged at 2000 g for 2 min, and then transferred into the well plate (∼500 µL per well). The granular hydrogel groups with the addition of fibrinogen in the well plate were incubated at ∼37℃ overnight before measurement. Bulk hydrogel controls were formed and crosslinked inside the well plate: 80 mg mL^−1^ GelMA in PBS with 2.5 mg mL^−1^ LAP was crosslinked using a UV lamp (365 nm, ∼6.8 mW cm^−2^, Spectroline ENF-260C/F) for 10 min and bulk fibrin gels were formed by mixing 490 µL of 10 mg mL^−1^ fibrinogen solution with 10 µL of 1 U mL^−1^ thrombin solution. Oscillatory stain sweeps were performed from 0.1-1000% strain at 10 rad s^−1^. The plate was lowered until the full surface of the gel was probed. The oscillatory frequency sweeps were run at 1-100 rad s^−1^ under 0.1% strain. Viscosity measurements were measured over a rotational shear rate of 0.01-100 s^−1^. Shear ramps tests were set to measure the shear moduli under alternating low strain (0.1% strain) and high strain (200%) at a constant frequency of 10 rad s^−1^ (2.5 cycles, 110 s per cycle). Time sweeps were measured overnight at constant frequency (10 rad s^−1^) and strain (0.1%).

### Compression Testing using a Dynamic Mechanical Analyzer

Dynamic cyclic compression testing was performed using a DMA850 (TA Instruments) fitted with a submersion compression clamp at 37℃. MG and Thr-MG in complete endothelial cell media were jammed into granular hydrogels (GH, Thr-GH), and loaded into a 2.5 mL Terumo Syringe. ∼80 µL of the granular hydrogel samples were extruded into separate cylindrical molds with a diameter of 6 mm and height of 3 mm. 20 µL of 10 mg mL^−1^ fibrinogen in endothelial cell medium was added to the granular hydrogels and samples were incubated overnight at 37℃. Bulk GelMA controls were prepared using a similar protocol, with 80 mg mL^−1^ GelMA precursor and 2.5 mg mL^−1^ LAP in complete endothelial cell media added to the mold, crosslinked using UV light (∼6.8 mW cm^− 2^, 10 min). Calipers were used to measure the dimensions of the samples before they were loaded onto the testing rig and secured with a clamp at a preload of 0.01 N. The chamber was then filled with ∼4 mL complete endothelial cell media to fully immerse the sample. Oscillation strain sweeps were performed after a 3-min soaking period to reach equilibrium under force control with 1% strain increments at 1 Hz and a force track of 115%. The force track feature was enabled to maintain the ratio between static and dynamic forces, ensuring that static force scaled with the stiffness of the samples. This adjustment prevented overstressing samples during cyclic loading. While the force track feature facilitated dynamic adjustments for consistent amplitude without overstraining, it did not directly control the thickness or modulus. The thickness of the samples was also tracked during the tests. Initially, a sweep from 0.01-60% strain was conducted to determine the failure strain of each sample. Subsequent sweeps were performed from 0.01-25% for GelMA and Thr-GH + Fib, and from 0.01-15% for GH + Fib (these samples could not reach 25% strain). Dynamic compressive moduli were calculated using the gradient of the stress-strain plot between 2-7% strain. To evaluate the cyclic performance of Thr-GH + Fib, the sweeps from 0.01-25% strain were performed for 30 cycles.

### Cell Culture

Human umbilical vein endothelial cells (HUVECs, Merck SCCE001) were cultured using complete endothelial cell media consisting of 5% v/v fetal bovine serum, 1% v/v endothelial cell growth supplement and 1% v/v penicillin-streptomycin in endothelial cell basal medium (Caltag Medsystems SC-1001). All cells were cultured in T-75 flasks (Greiner Bio-One 658175) coated with fibronectin (Merck F1056) at ∼2 µg cm^2^, with media changed every two days and passaged at a confluency of 80% using 0.05% v/v trypsin-EDTA (Merck T3924) in PBS. All cells were used at passage nine or lower. All microgels used for cell culture were sterilized under UV (254 nm, ∼0.7 mW cm^−2^, Spectroline ENF-260C/FE) for 1 h, while the thrombin functionalization was performed as described above but under sterile conditions. All chemical reagents were either sterile-filtered (GelMA, LAP, EDC, sulfo-NHS, MES buffer, fibrinogen), autoclaved (agarose), or prepared under sterile conditions (thrombin, PBS).

### Imaging Cell Attachment and Proliferation

10 mg mL^−1^ ultrapure agarose in PBS (Fisher Scientific 16550100) was prepared, sterilized, and reheated in a microwave (HADEN 193926) for 60 s. 500 µL of warm agarose solution was added to separate wells of a 48-well plate and gelled at 4℃ for 1 h. This was to ensure that the samples were close to the surface of the well to be imaged under upright multiphoton microscope. Sterile, rhodamine-tagged microgels (MG or Thr-MG) were jammed into granular hydrogels on top of the agarose gel in the 48-well plate (∼150 µL per well). The granular hydrogels were then incubated overnight in either 50 µL of 10 mg mL^−1^ fibrinogen in complete endothelial cell media (for Thr-GH + Fib, GH + Fib) or complete endothelial cell media alone (Thr-GH). A suspension of HUVECs at 1.7 × 10^5^ cells mL^−1^ was prepared in complete endothelial cell media and 300 µL was seeded onto each sample. Half media changes (150 µL) were performed every 2 days, with samples harvested after 1, 4, or 7 d of culture. These samples were fixed using 40 mg mL^−1^ formaldehyde (Merck P6148) in PBS for 15 min and then washed 3 times with PBS. The samples were permeabilized in 0.1% v/v Triton X-100 in PBS for 30 min, blocked with 0.3% v/v Triton x-100 and 5 mg mL^−1^ BSA in PBS for 30 min. The samples were stained through incubation with a 660 nM solution of Alexa Fluor 488 phalloidin (Thermo Fisher Scientific A12379) for 2 h and a 1 µg mL^−1^ solution of 4’,6-diamidino-2-phenylindole dihydrochloride (DAPI, Thermo Fisher Scientific 62248) for 30 min. The samples were then washed 3 times with PBS for 10 min after each step. The stained samples were kept at 4℃ before imaging. Stained samples were imaged using a Leica SP8 AOBS confocal laser scanning microscope attached to a Leica DM6000 upright epifluorescence microscope (multiphoton microscope). Image stacks were acquired with a step size of 3.77 µm using a 10X dry objective lens, a 488 nm argon laser, a 561 nm solid state yellow laser and a 750 nm multiphoton laser. Acquired images were processed using Fiji.

### Quantifying Cell Metabolism

An alamarBlue assay (Thermo Fisher Scientific DAL1025) was used to quantify the metabolic activity of the seeded cells over time (n = 3). Samples were prepared as described for the imaging study using non-fluorescent microgels and 2 × 10^4^ cells per well. After 1, 4, and 7 d in culture, 300 µL of 20% v/v alamarBlue reagent in complete endothelial cell media was added to each well to yield a final alamarBlue reagent concentration of 10% v/v. The samples were incubated at 37℃ for 1 h. 400 µL of supernatant was taken from each well and centrifuged in an Eppendorf tube to remove any unbound microgels or cells. 100 µL triplicates of each supernatant were then transferred to a black 96-well plate. Fluorescence readings were measured (λ_ex_ = 570 nm, λ_em_ = 585 nm) using a microplate reader. The supernatant was replaced with 300 µL fresh cell media, and half media changes (150 µL) were performed every 2 days.

### Vascular Spheroid Invasion Assay

HUVECs and hMSCs were harvested from culture flasks with trypsin and counted using an automatic cell counter (Denovix CellDrop FL). The cells were combined at a 2:1 ratio and resuspended in a 2:1 mixture of HUVEC media and hMSCs media to a total cell concentration of 1 × 10^4^ cells mL^−1^. This cell suspension was transferred into ultra-low attachment 96-well plates (Merck 7007) with 2000 cells per well. The well plate was tapped gently and cultured for 3 days to form vascular spheroids. These spheroids were then cultured with the biomaterial (Thr-GH or Thr-GH + Fib) as previously reported. ^[34]^ Briefly, the spheroids were harvested into Eppendorf tubes and then transferred to a black 96-well plate pretreated with anti-adherence rinsing solution (Stem Cell Technologies 07010) (∼40 spheroids per well). Sterile Thr-GHs were transferred into the 96-well plate (∼100 µL per well) and gently mixed with spheroids using a pipette tip. 20 µL of 10 mg mL^−1^ fibrinogen in the 2:1 mixed cell media was added to half of the wells to form Thr-GH + Fib composite. After 4 h of incubation, 100 µL of the 2:1 mixed media was added to each well. After 1 d of culture, the media was replaced with 100 µL of fresh 2:1 mixed media supplemented with 100 ng mL^−1^ recombinant human vascular endothelial growth factor (Thermo Fisher Scientific PHC9394). The samples were cultured for another 2 days before fixation with 40 mg mL^−1^ formaldehyde in PBS for 40 min and then washed 3 times with PBS. The samples were then permeabilized, blocked, and stained with Alexa Fluor 488 phalloidin and DAPI, as described above, except that phalloidin was stained through incubation overnight at 4℃. Stained samples were imaged using a Leica SP8 multi-laser confocal laser scanning microscope equipped with a 488 nm argon laser and 561 nm solid state yellow laser. Images were acquired using 10X dry objective lenses with a step size of 4.28 µm used for 3D stacks. Acquired images were processed using Fiji. Horizontal outgrowth was calculated by normalizing the average radii of the day 3 vascular networks from maximum projection images to the radius of a day 0 spheroid. Vertical outgrowth was calculated using the radius of a day 0 spheroid (r_0_) to normalize the vertical displacement in the day 3 vascular networks (total depth – r_0_).

### Statistical analysis

Statistical analysis and plotting of rheological experiments and compression tests were performed using Origin 2024b (OriginLab). Statistical analysis and plotting of biological experiments were performed using GraphPad Prism 10. Shapiro-Wilk normality test was performed and statistical significance was assessed using Kruskal-Wallis test with Dunn’s post-hoc test, unpaired t test with Welch’s correction, and two-way ANOVA with Tukey post-hoc test.

## Acknowledgements

The authors acknowledge the Wolfson Bioimaging Facility for their support and assistance in microscopy. Z.D. acknowledges the Tissue and Cell Engineering Society (TCES) for the Adam Curtis Collaboration Award and support from the China Scholarship Council – University of Bristol PhD Scholarship (CSC202201130001). S.B. thanks the University of Glasgow for funding. F.S. and G.C. acknowledge the support from the ERC-2020-AdG 101020715 NEUROMETA project. M.R.W. acknowledges the support from the NIHR Biomedical Research Centre at University Hospitals Bristol and Weston NHS Foundation Trust and the University of Bristol. The views expressed are those of the authors and not necessarily those of the NIHR or the Department of Health and Social Care. J.P.K.A. acknowledges funding from a UKRI Future Leaders Fellowship (MR/V024965/1). J.P.K.A. and M.R.W. acknowledge the EPSRC project “emPOWER: in-body artificial muscles for physical augmentation, function restoration, patient empowerment and future healthcare” (EP/T020792/1). Schematics used in figures were created with Biorender.com.

## Table of Contents

*Coagulative granular hydrogels* are composed of packed thrombin-functionalized microgels that catalyze the conversion of fibrinogen into a secondary fibrin network, filling the interstitial voids. This bio-inspired approach stabilizes the biomaterial to match the robustness of bulk hydrogels without compromising injectability, mimicking the initial stage of wound healing and providing a conducive microenvironment for cell adhesion and proliferation.

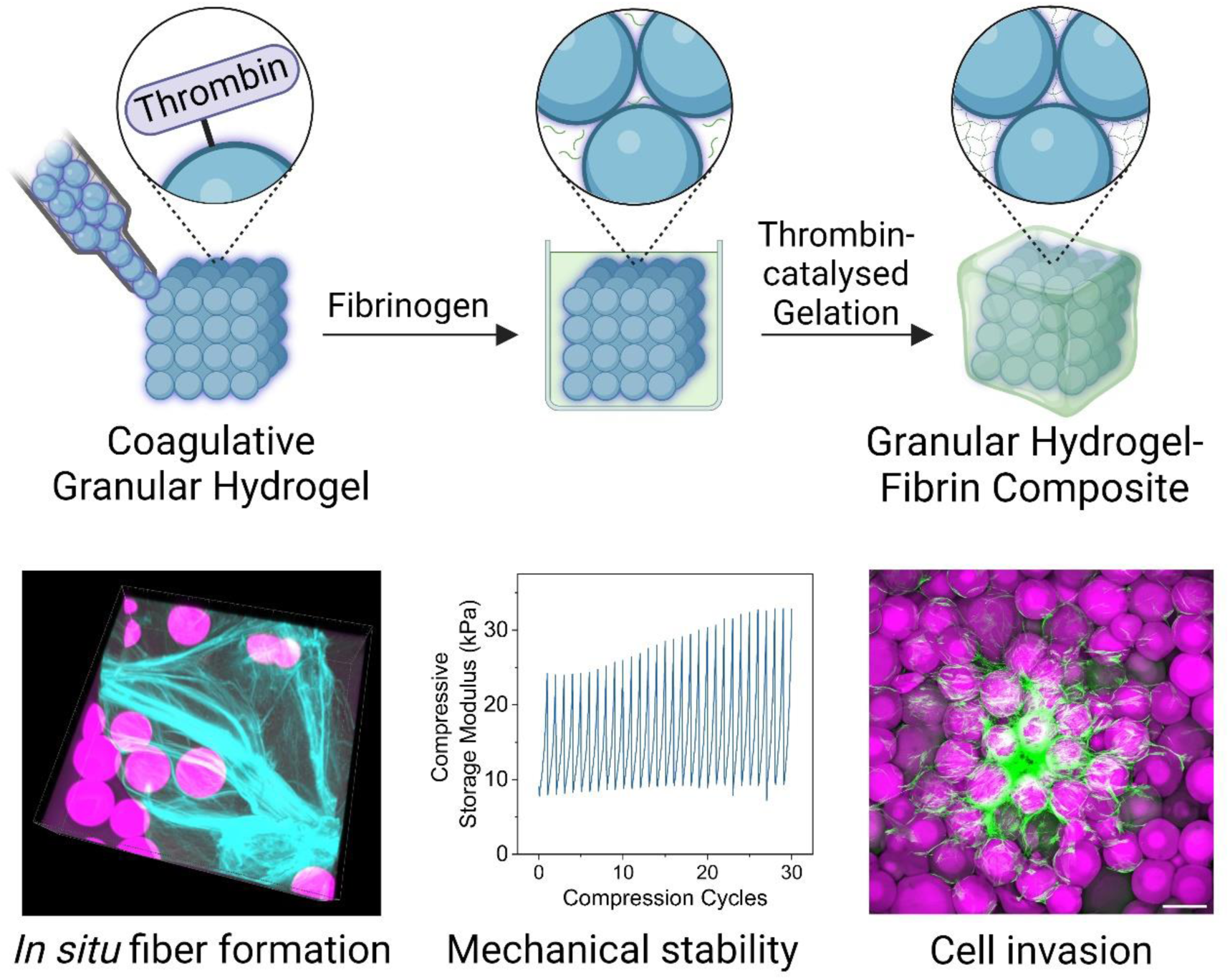

## Notes

### Competing Interest Statement

The authors have declared no competing interest.

