## Supporting information for "Coagulative Granular Hydrogels with an Enzyme Catalyzed Fibrin Network for Endogenous Tissue Regeneration"

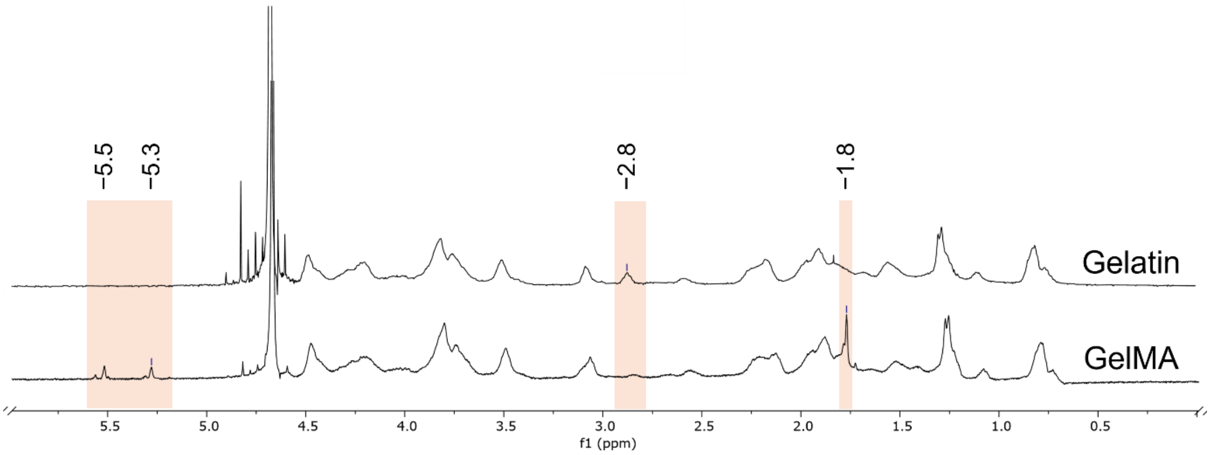


**Figure S1**. Representative $^1$H NMR spectra of GelMA and gelatin. The spectrum of GelMA displayed peaks corresponding to acrylic protons (2H) of methacrylamide on lysine sidechains (~5.3 ppm) and hydroxylysine sidechains (~5.5 ppm), and methyl protons (3H) of methacrylamide on either lysine or hydroxylysine sidechains (~1.8 ppm). These three peaks, which were not present in gelatin, indicated a successful conjugation of methacryloyl groups. The near absence a methylene lysine proton peak (2H) (~2.8 ppm), which was present in the spectrum of gelatin, indicated that the free amino groups of gelatin were efficiently reacted.[1]

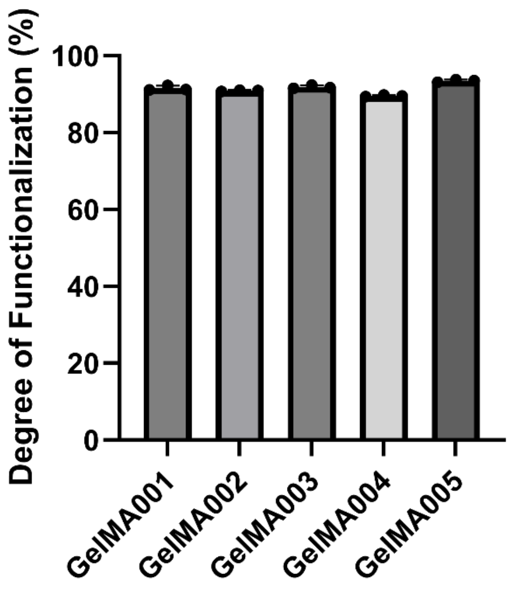


**Figure S2**. The degree of functionalization of the five GelMA batches (labelled 001-005) used in this study, which was calculated from a fluoraldehyde assay against gelatin standards. Data shown as mean ± standard deviation, n = 3.

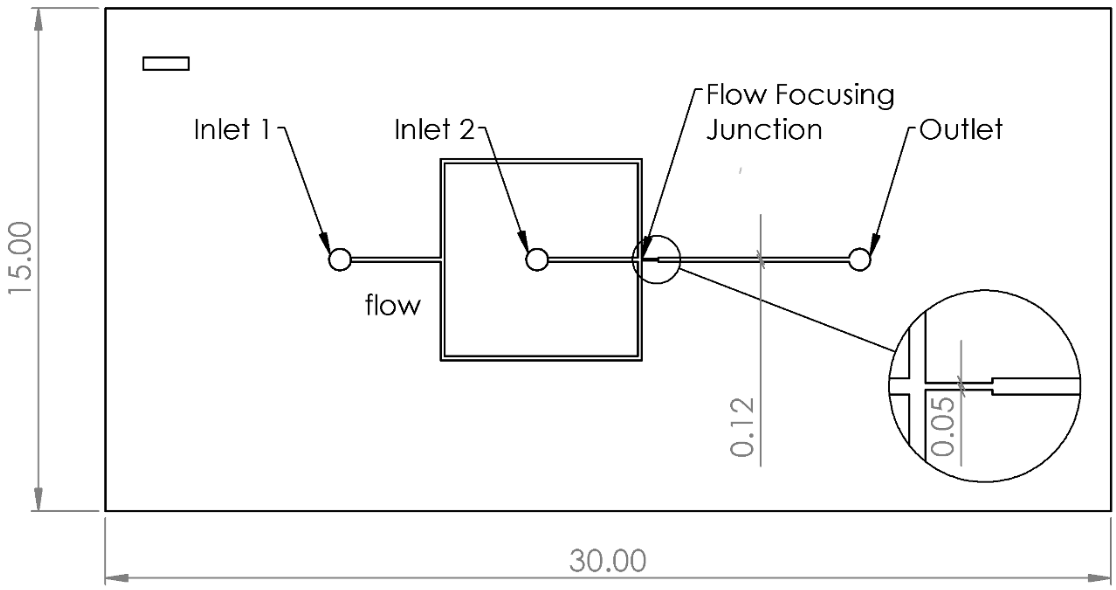


**Figure S3**. Design of the microfluidic chip for microgel fabrication. The surfactant-supplemented oil phase enters via inlet 1 and the liquid hydrogel precursor phase enters via inlet 2. The two liquids meet at the flow focusing junction, which generates hydrogel precursor microdroplets in a continuous flow of oil. The microgel droplets exit via the outlet and are then crosslinked with UV radiation. The channel has a constant width of 0.12 mm, except at the orifice of the flow focusing junction, where it narrows to 0.05 mm. Units in mm.

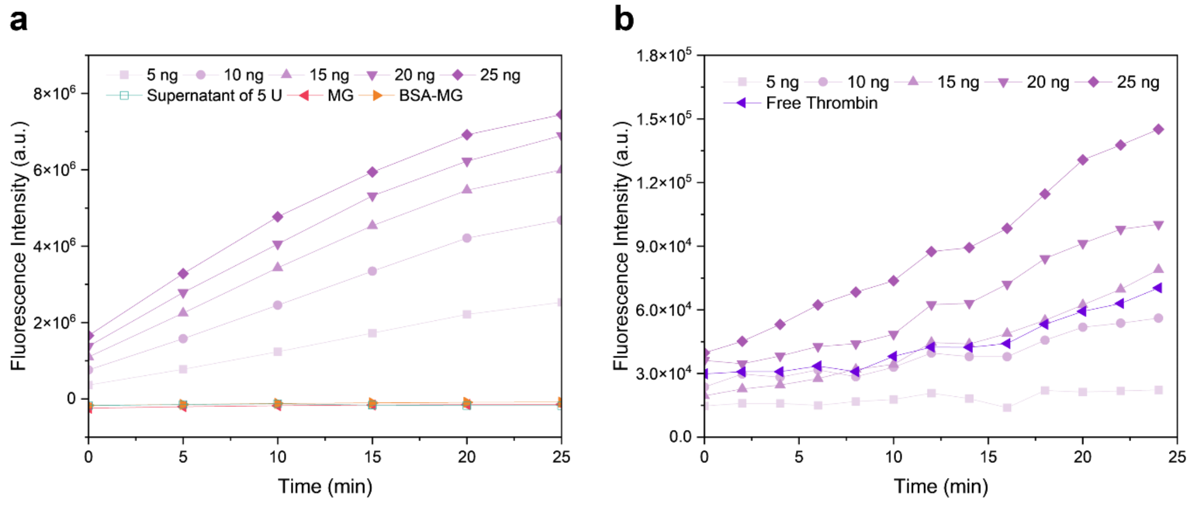


**Figure S4**. Thrombin activity assay. a) Activity assay curves of all controls: unfunctionalized microgels, BSA-functionalized microgels, and the supernatant from a 5 U thrombin functionalization reaction. When compared to the thrombin standards, these controls all showed negligible enzyme activity. b) Activity assay curve of thrombin reagent solution incubated overnight against fresh thrombin standards. The calculated activity is ~10 ng, which is 43% of the activity (23 ng) of fresh thrombin at the same concentration.

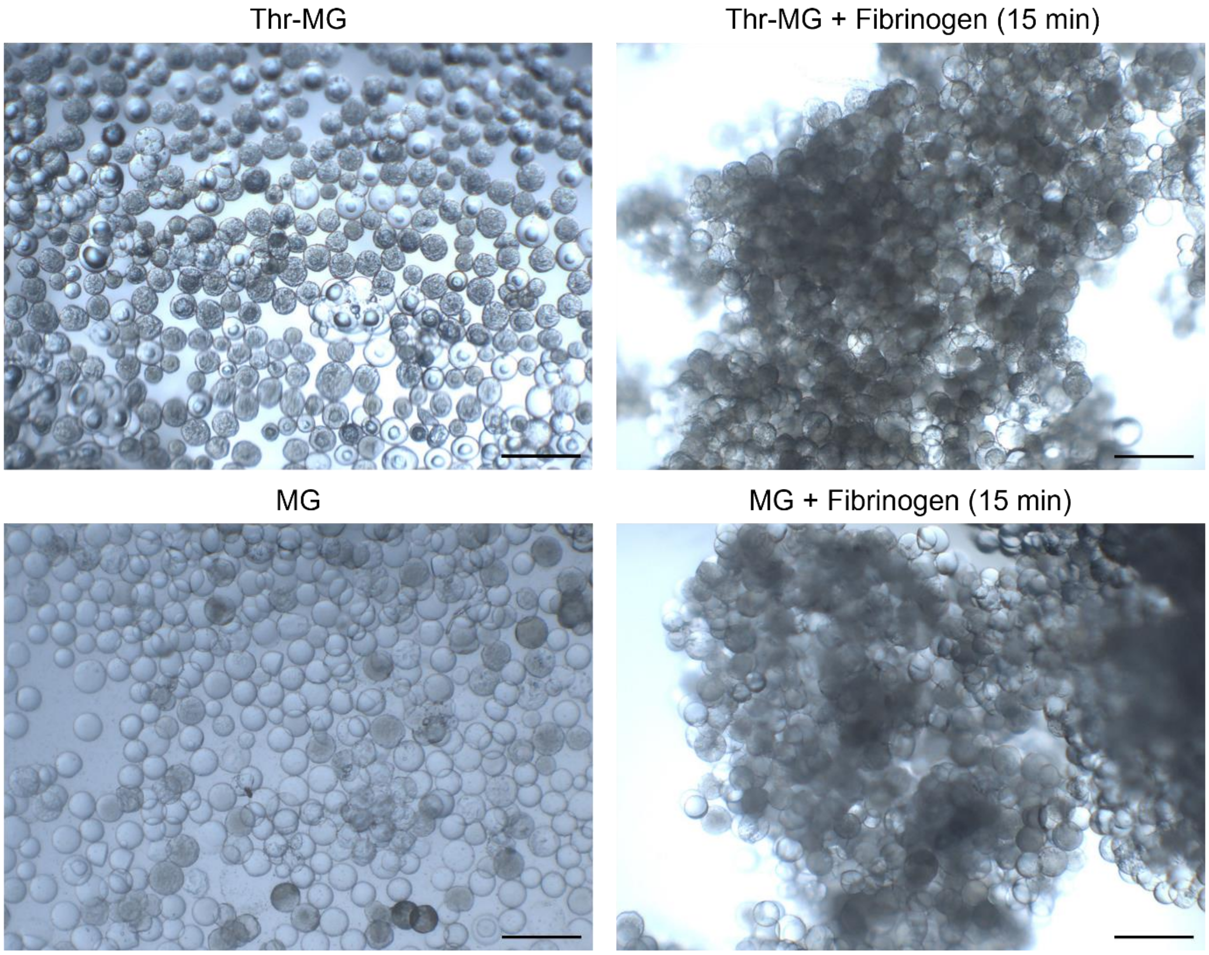


**Figure S5**. Optical images of Thr-MG and unfunctionalized microgels (MG) without any fibrinogen and 15 minutes after the addition of fibrinogen. Aggregation of microgels was observed in both groups at the early timepoint. Scale bar: 200 µm.

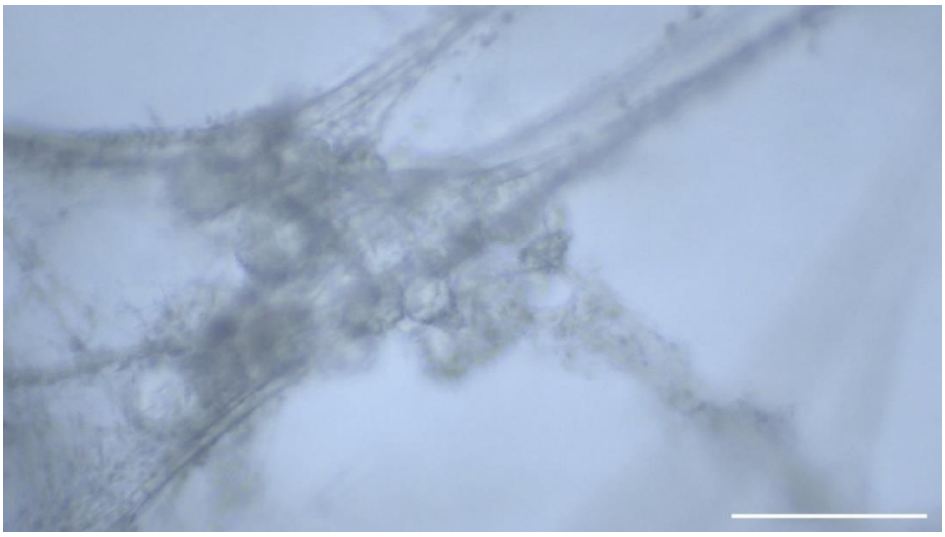

**Figure S6**. Brightfield image of Thr-MG aggregates incubated with fibrinogen for 24 h showed the presence of fibers, assumed to be fibrin, entangled with the Thr-MGs. Scale bar: 200 µm.

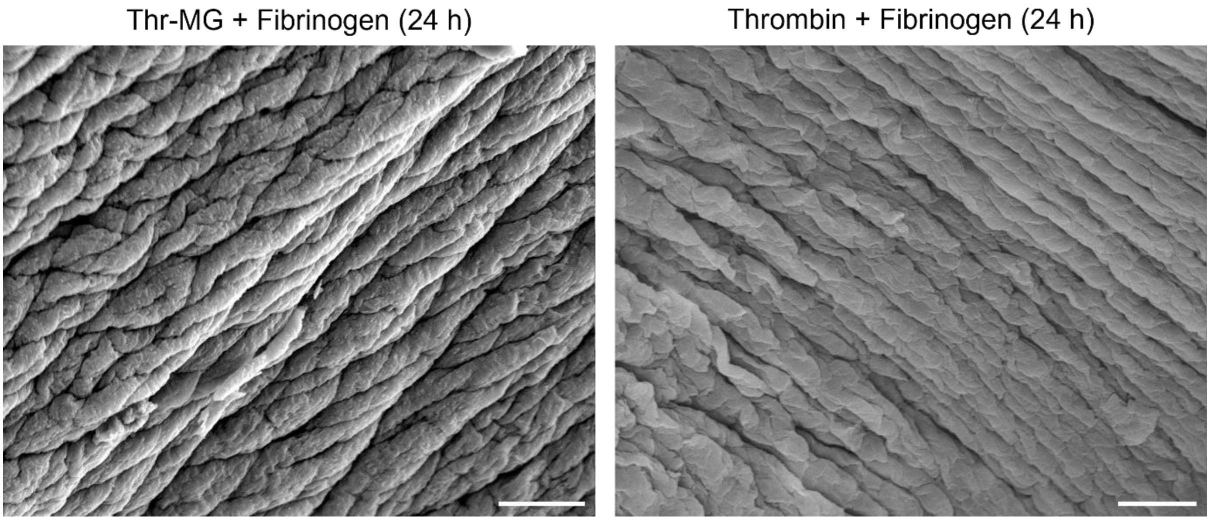


**Figure S7**. Representative scanning electron microscopy images of fibrin formed by Thr-MG (left) and thrombin solution (right). These samples showed similar morphology, with both exhibiting fibrillar structure and hierarchical organization. Scale bar: 10 µm.

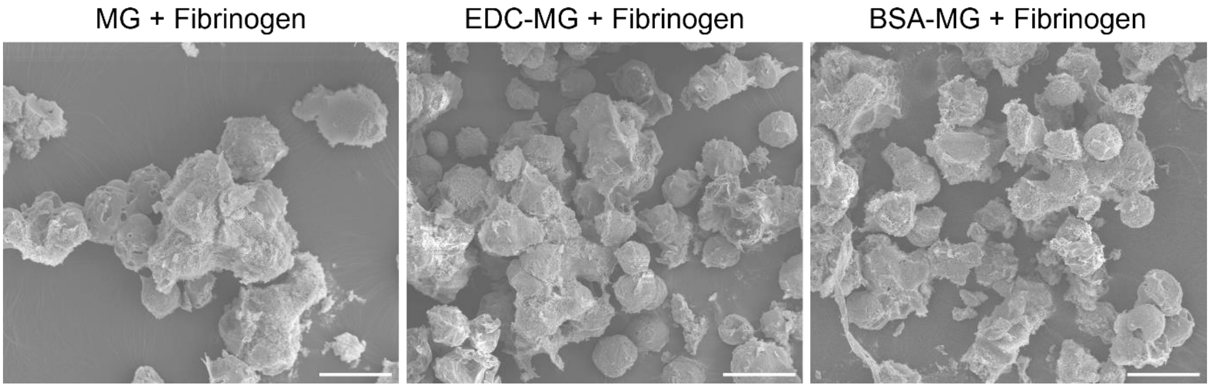


**Figure S8**. Representative scanning electron microscopy images of microgels (MG), EDC-activated microgels (EDC-MG), BSA-functionalized microgels (BSA-MG), each incubated with fibrinogen for 24 h. Scale bar: 100 µm.

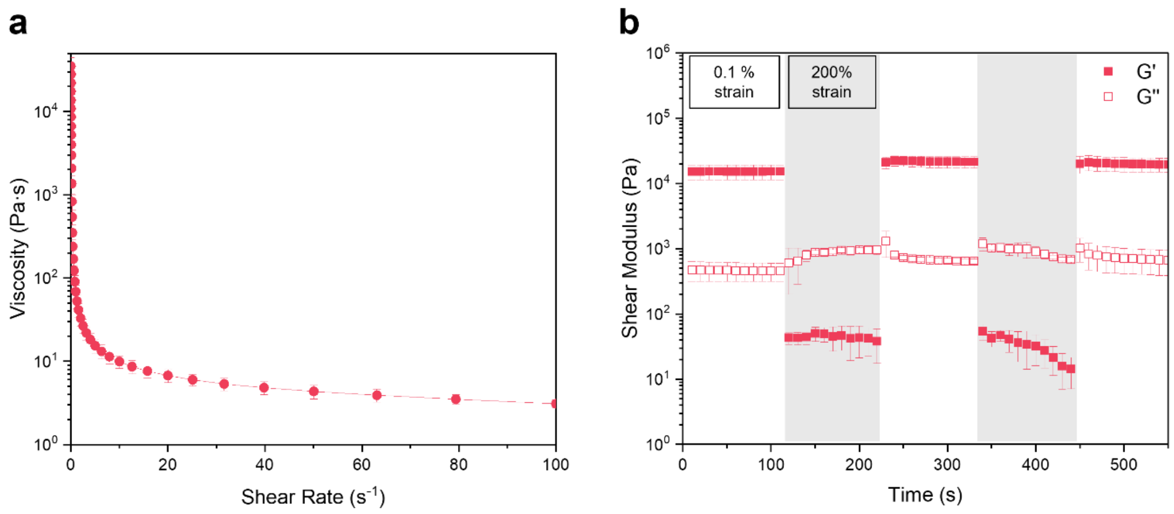


**Figure S9**. a) Viscosity measurement of unfunctionalized granular hydrogels (GH) with shear rates from 0-100 $s^{-1}$. b) Shear recovery test of GH, measuring shear storage modulus (G') and shear loss modulus (G") when alternating between 0.1% strain (unshaded) and 200% strain (shaded). Data shown as mean ± standard deviation, n = 3.

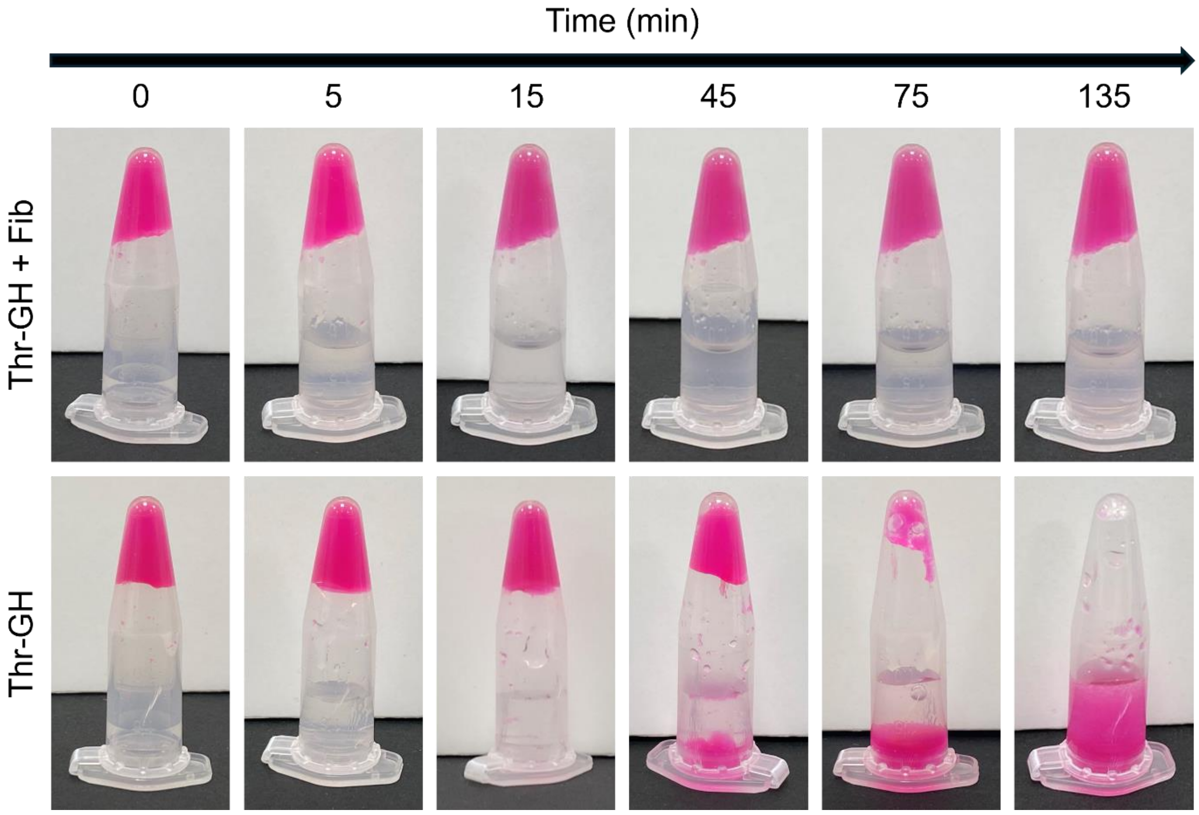


**Figure S10**. The stability of Thr-GH + Fib and Thr-GH control was assessed by exposing the samples to continuous agitation. The Thr-GH + Fib composite remained intact under the dynamic conditions, while the Thr-GH control began to disintegrate after 15 min of agitation and was completely dissociated after 2 h.

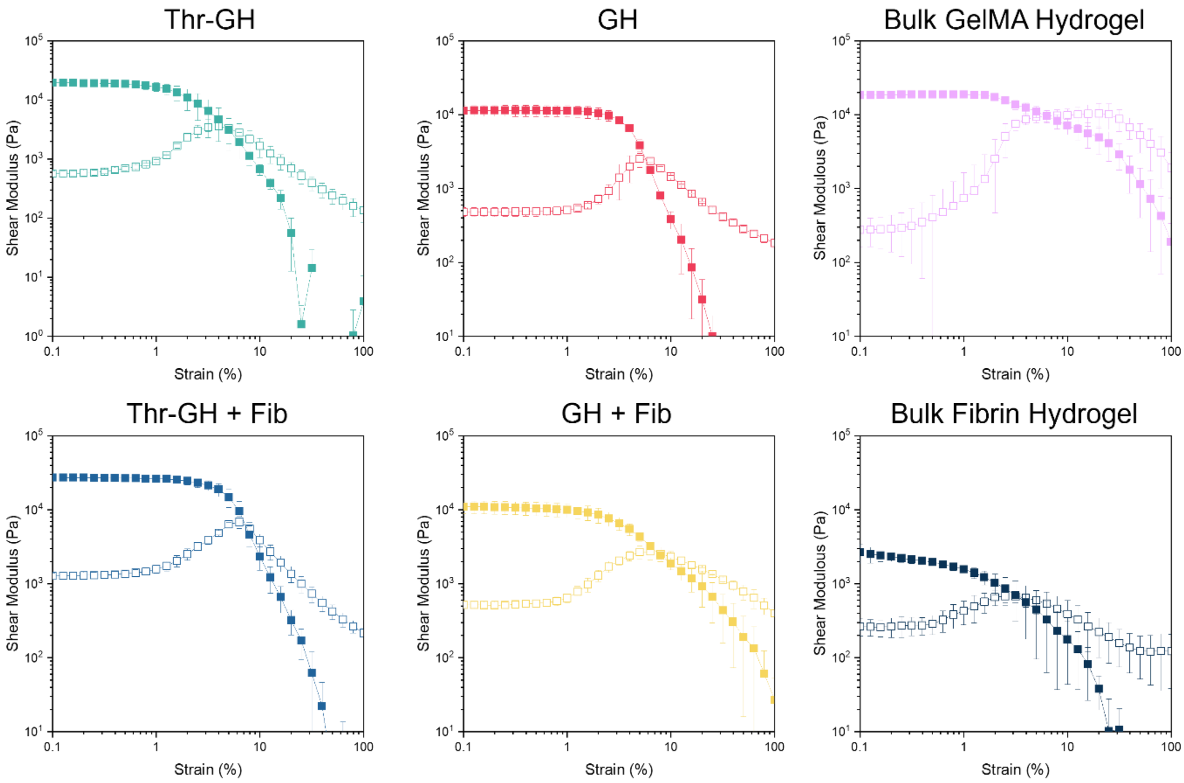


**Figure S11**. Oscillatory shear strain sweeps of Thr-GH, Thr-GH-fibrin composite (Thr-GH + Fib), GH, GH with the addition of fibrinogen (GH + Fib), bulk GelMA hydrogel, and bulk fibrin hydrogel. The shear storage modulus (G', filled symbols) and shear loss modulus (G'', open symbols) were measured across a shear strain range of 0.1-100%. Data shown as mean ± standard deviation, n = 3.

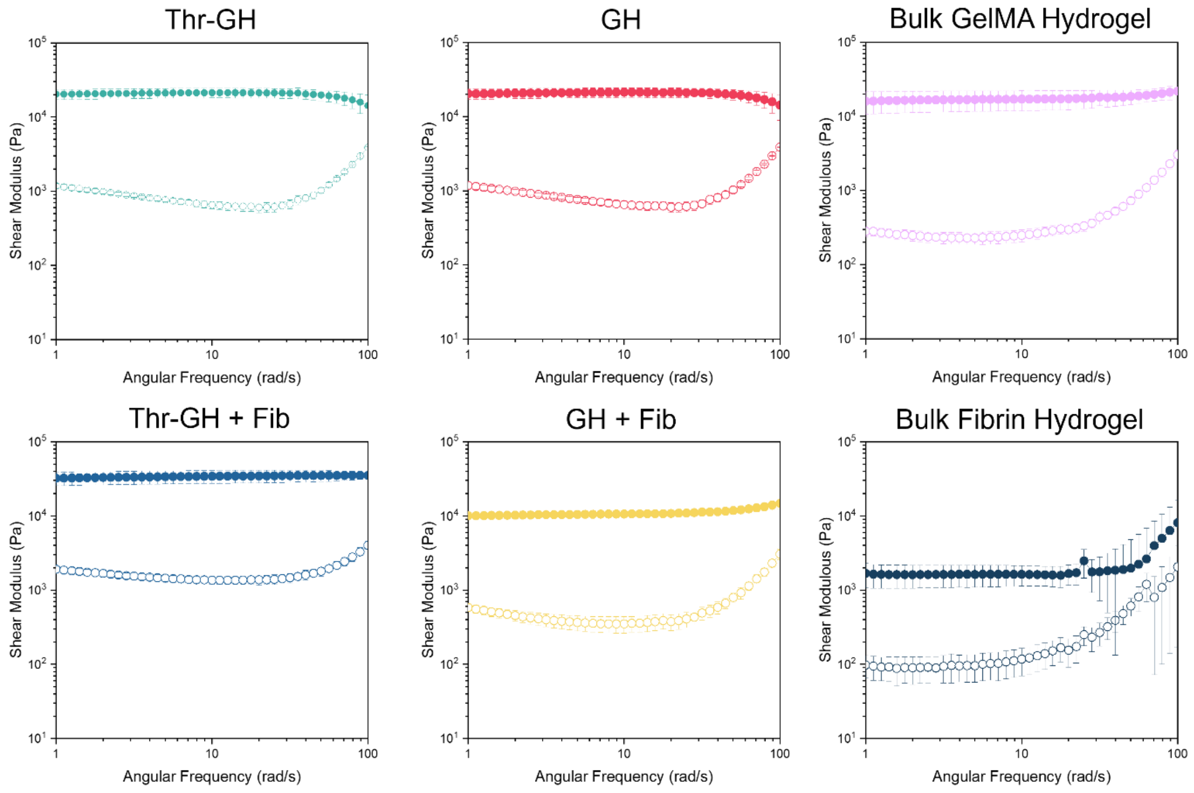


**Figure S12**. Oscillatory shear frequency sweeps of Thr-GH, Thr-GH + Fib, GH, GH + Fib, bulk GelMA hydrogel, and bulk fibrin hydrogel. The shear storage modulus (G', filled symbols) and shear loss modulus (G", open symbols) were measured across an angular frequency range of 1-100 rad/s. Data shown as mean ± standard deviation, n = 3.

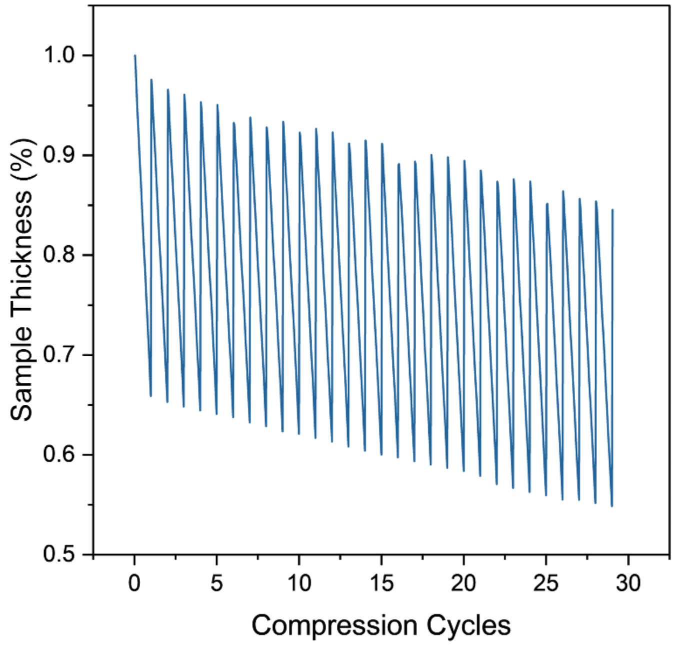


**Figure S13**. The thickness recovery of Thr-GH + Fib during dynamic compression.

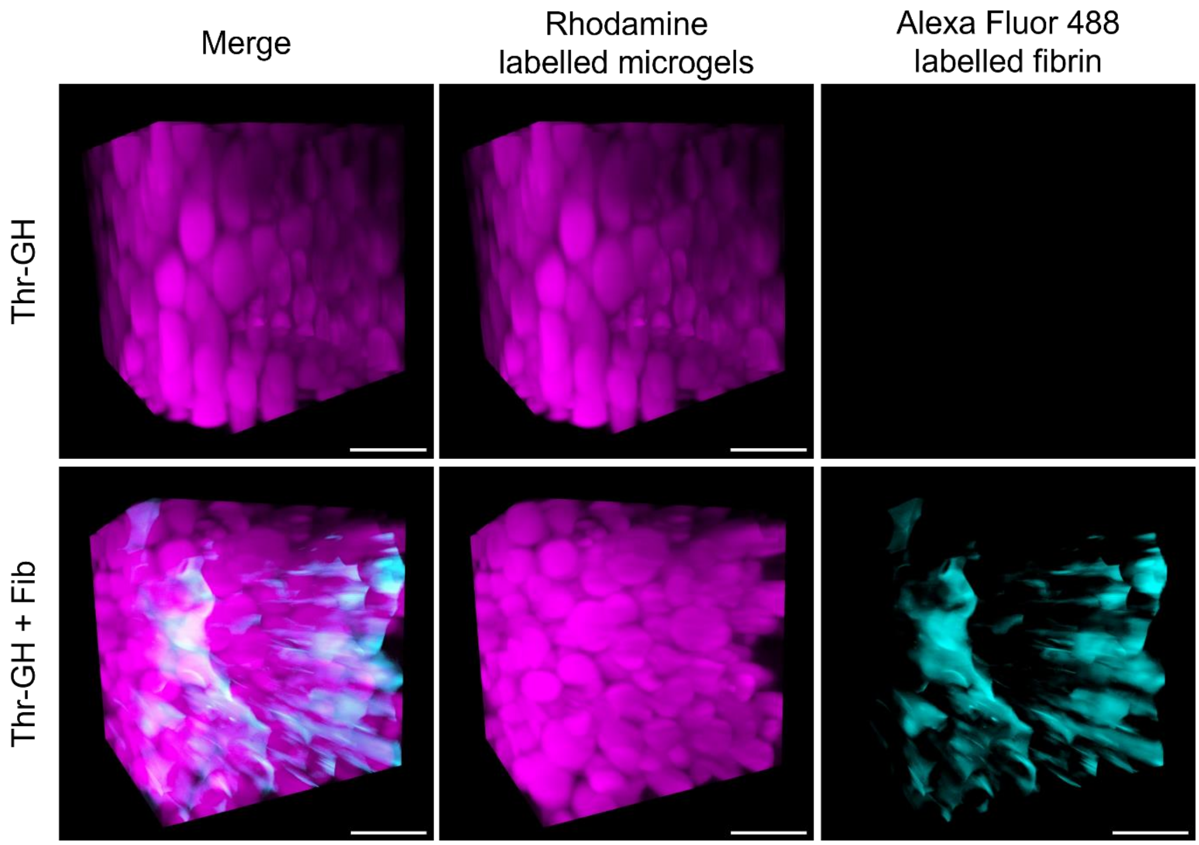


**Figure S14**. 3D reconstructed stacks of Thr-GH (top panel) and Thr-GH + Fib (bottom panel) from light sheet fluorescence microscopy. Microgels were made with GelMA fluorescently labelled with rhodamine (shown in purple), while the Thr-GH sample was incubated with Alexa Fluor 488 labelled fibrinogen (shown in cyan). Scale bar: 500 µm.

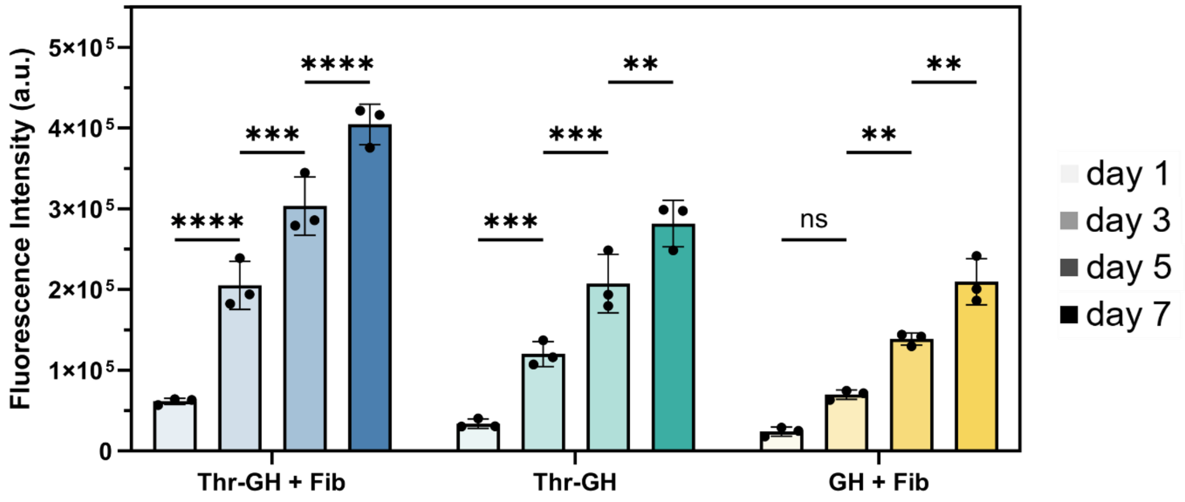


**Figure S15**. Quantification of cell metabolism in Thr-GH + Fib, Thr-GH and GH + Fib over 7 days. Data are presented as mean ± standard deviation, with a sample size of n = 3. Statistical analysis performed using two-way ANOVA with Tukey's post-hoc test; ns = no significance ($p > 0.05$), **$p < 0.01$, ***$p < 0.001$, ****$p < 0.0001$.
